# Large-area electrical imaging having single neuron resolution using 236,880 electrodes CMOS-MEA technology

**DOI:** 10.1101/2022.11.09.515884

**Authors:** I. Suzuki, N. Matsuda, X. Han, S. Noji, M. Shibata, N. Nagafuku, Y. Ishibashi

## Abstract

The electrophysiological technology having a high spatio-temporal resolution at the single-cell level, and noninvasive measurements of large areas provides insights on underlying neuronal function. Here, we used a complementary metal-oxide semiconductor (CMOS)-microelectrode array (MEA) that uses 236,880 electrodes each with an electrode size of 11.22 × 11.22 µm and 236,880 covering a wide area of 5.5 × 5.7 mm in presenting a detailed and single-cell-level neural activity analysis platform for brain slices, human iPS cell-derived cortical networks, peripheral neurons, and human brain organoids. Propagation pattern characteristics between brain regions changes the synaptic strength into compounds based on single-cell time-series patterns, classification based on single DRG neuron firing patterns and compound responses, axonal conduction characteristics and changes to anticancer drugs, and network activities and transition to compounds in brain organoids were extracted. This detailed analysis of neural activity at the single-cell level using our CMOS-MEA provides a new understanding the basic mechanisms of brain circuits *in vitro* and *ex vivo*, on human neurological diseases for drug discovery, and compound toxicity assessment.

## Introduction

The function of the nervous system is represented by an electrical activity. The technology for measuring the electrical activity of the nervous system is essential for understanding higher functions and neurological diseases, drug discovery development, and toxicity evaluation of compounds. The methods for measuring electrical activity include electrical measurement, optical measurement, and magnetic field measurement. Electrical measurement has a high temporal resolution, and optical measurement has a high spatial resolution. In the recent years, the density of microelectrode array (MEA) has increased, and the spatial resolution in electrical activity measurement has increased as well. With the latest advances in complementary metal-oxide semiconductor (CMOS) technology, recent large-scale neural recording technologies are significantly increasing the number of single cells/units which can be recorded simultaneously at multiple sites of the brain^1,2,3,4^. CMOS-based shank silicon probes cover the important brain regions including the hippocampus, cortex, and thalamus, observed in rats and mice, to provide long-term stable recording after chronic implant^5,555, 6, 7, 89^, 10, 11, 12. With such CMOS-based probes, extracellular action potentials from around 1,000 individual single neurons could be monitored simultaneously from different brain structures, providing global insight into neural activities to understand neural network function in animal brains^13^. Using MEA for *in vitro* and *ex vivo*, CMOS-MEA development has progressed. High-density (HD) CMOS-MA capable of simultaneous recording signal from approximately 1,000 electrodes to a maximum of 19,000 electrodes has been developed^14^. Axonal conduction measurements in cultured neurons and electrical activity patterns in brain slices and retinal tissue have been reported^15^. The development of electrical imaging by HD-CMOS-MEA makes it possible to link the structure and electrical activity of neural circuits at the single-cell level. The demand for *in vitro* electrical activity measurement of the nervous system using MEA has increased during the recent years. The discovery of human iPS cells (hiPSC)^16^ has made it possible to use human neurons. This is because the elucidation of human neurological diseases and the development of new drugs will advance from extrapolability to humans and the use of human neurons derived from disease patients. The toxicity evaluation of pharmaceuticals using healthy hiPSC-derived neurons^15, 17, 17, 18, 19, 20, 21, 22, 23, 24, 25, 26, 27, 28, 29, 30, 31^ and the electrical activity analysis of neurons from diseased patients has been reported^32, 33, 34, 35, 36, 37, 38, 39, 40^. Additionally, hiPSC-derived peripheral neurons are used in evaluating pain associated with pain-related substances and anticancer drug administration compounds^41, 42, 43, 44, 45, 46^. Some brain organoids which mimic the three-dimensional structure of humans have been studied with MEA measurement method used as a functional measurement method for brain organoids^47, 48, 49, 50, 51, 52^. HD-CMOS-MEA captures activity and axonal conduction at the single-cell level because of its high spatial-temporal resolution. By measuring activity at the single-cell level, it will be possible to quantify propagation patterns via synapses in the neuronal networks and cerebral organoids using extracellular potentials. This will contribute to *in vitro* drug efficacy assay and neurological disease research. Peripheral nerves which control pain are known to have different channels expressed in various that have different cells responses to compounds^41, 53, 54, 55, 56^. Additionally, the peripheral neuropathy by the compound is divided into toxicity to somas, axons, and myelin. By evaluating pain-related substances using the firing pattern of a single cell, it will become a new pain evaluation system based on the electrical activity. Additionally, when combined with axonal conduction measurements, it will become an assessment method to predict soma, axonal, and myelin toxicity based on electrical activity.

HD-CMOS-MEA is a technology that enables the measurement of nerve activity and axonal conduction at the single-cell level. However, to analyze the neural network activity state *in vitro* and *ex vivo*, it is a challenge to develop a measurement system which combines the following five points: (1) high spatial resolution, (2) high temporal resolution, (3) wide range measurement, (4) high signal to noise ratio (SNR), and (5) real-time measurement with fast scan speed. Our HD-CMOS-MEA features 236,880 platinum (Pt) electrodes with a size 11.22 × 11.22 μm and pitch of 0.25 μm, and 33,840 readout channels operating at 70 kSamples/second (kS/s) with a noise level of 5.5 μVrms. In addition, our MEA enables simultaneous measurement of a wide area of 5.51 × 5.91 mm^2^ (Figure 1). This MEA employs a disaggregated differential amplifier and one-sided feedback and auto-zero circuit to reduce the pixel size and suppress the noise. Moreover, it integrates single-slope ADCs, a 4.752-Gbps/ch output interface, and a stacked device structure to enhance the readout speed^57^. Due to densely arranged electrodes, single-cell signals can be recorded with multiple electrodes. The noise is sufficiently suppressed by processing an average or a correlation between the signals from these electrodes, and small signals, such as axonal signals or action potentials of immature neurons, can be detected. The large sensing area can visualize the entire neuronal network activities while recording signals at a cellular resolution.

**Figure 1.**
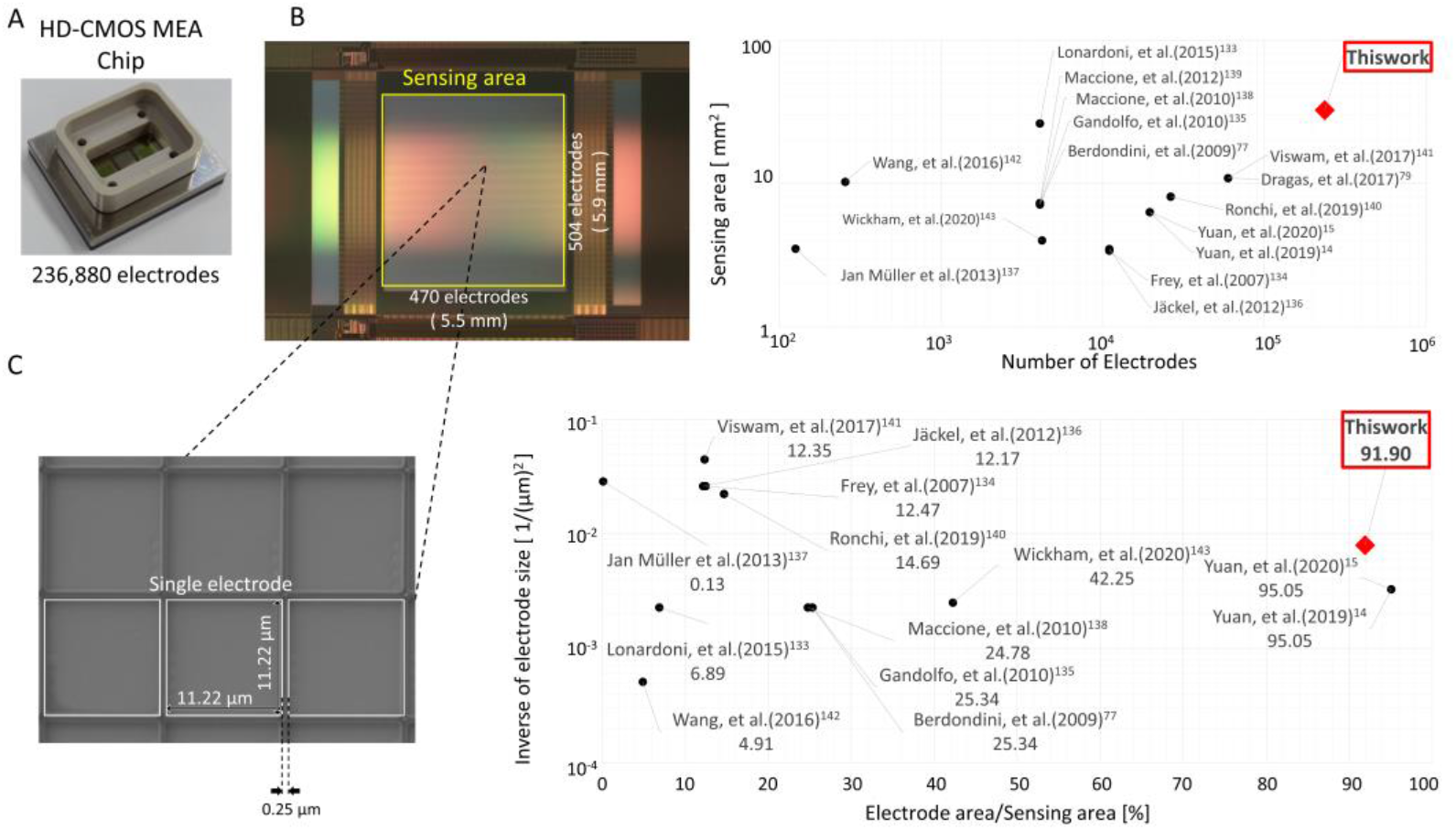
High-density (HD) CMOS-MEA with 236,880-electrode and wide measurement area. (A) Overview of HD-CMOS-MEA chip. (B) Sensing area (5.5 mmH × 5.9 mmV). (C) Electron microscope image of single electrode size (11.22 µm × 11.22 µm, pitch=0.25 µm). (D) Comparison of the number of electrodes to the measurement area^14, 15, 77, 79, 133–143^. (E) Comparison of the reciprocal of the electrode size to the number of electrodes per measurement area^14, 15, 77, 133–138, 140–143^.

In this study, using our HD-CMOS-MEA, we demonstrated (1) detection of detailed propagation patterns between regions in brain slices, (2) quantified synaptic strength and drug response based on single-cell-level firing patterns in human iPS central neural networks, and (3) detected of responses to pain-related substances based on the firing patterns of single-cell units in sensory neurons, and (4) detailed activity measurements and drug responses at the single-cell level in human cerebral organoids. These results are helpful for (1) understanding circuit operation principles based on propagation patterns obtained by detailed interregional propagation measurements in brain slices; (2) pharmacology based on single-cell-level firing patterns and synaptic strengths in the central neuronal network; (3) a novel assessment method for drug-induced peripheral neuropathy based on single cell-level firing patterns; and (4) neurological disease research and pharmacological tests based on propagation patterns of human cerebral organoids. These assays will provide a new and effective method for understanding neural circuit activity, including neurological diseases, and evaluating the efficacy and toxicity of pharmaceuticals, based on detailed propagation patterns of nerves *in vitro* and *ex vivo*.

## Results

### Performance of a 236,880-electrode CMOS-MEA and comparison with conventional CMOS-MEAs

Figure 1A–C shows the appearance of HD-CMOS-MEA with 236,880 platinum (Pt) electrodes, having a sensing area in 5.5 mmH × 5.9 mmV. It has an electron microscope image of the electrode surface with size 11.22 × 11.22 μm and pitch of 0.25 μm. The number of electrodes and sensing area is the largest compared to the conventional CMOS-MEAs (Figure 1D). The reciprocal value of the electrode area per electrode, that indicates the microscopic of the electrodes, is the fifth overall, but the electrode density to the measurement area is 91.9%. This is an HD-CMOS-MEA having the highest specs compared to other available MEA. It can measure nerve activity with a high spatial resolution because it uses microelectrodes and covers almost the entire sensing area (Figure 1E).

### Validation of large-scale and high spatial-temporal resolution recording capabilities on mouse brain slice

Our HD-CMOS-MEA with large-scale sensing area and high spatial resolution was used in recording extracellular activity in mouse sagittal brain slices. The HD-CMOS-MEA has sensing regions which allows simultaneous measurements from the cerebral cortex, hippocampus, midbrain, thalamus, and caudate putamen (Figure 2A), with spontaneous activity observed from the cerebral cortex and hippocampus (Figure 2B–D). Local field potential (LFP) reveals detailed propagation patterns for several hundred milliseconds in the hippocampus, perirhinal cortex (PC), and entorhinal cortex (EC) regions (Supplementary movie 1, Figure 2D). Figure 2C(a–f) shows the waveforms of 25 electrodes measured in 5 vertical and 5 horizontal electrodes (3147.21 µm^2^) in each region, which are indicated by the red dots in Figure 2B, and their average waveforms. The voltage values of the average waveform are 230 μV in cortical layer 5, 87 μV in cortical layer 3, 62 μV in cortical layer 1, 85 μV in the entorhinal cortex, 69 μV in the dentate gyrus, 525 μV in CA3, and 1622 μV in CA1. The signal-to-noise ratio (SNR) was 19.3 in cortical layer 5, 8.7 in cortical layer 3, 8.5 in cortical layer 1, 6.6 in the dentate gyrus (DG), 56.7 in CA3, and 136.2 in CA1 (Figure 1C– g). Although the signal intensity differed depending on the brain region, it was confirmed that the activity could be acquired with a high SNR from the spontaneously active region.

**Figure. 2.**
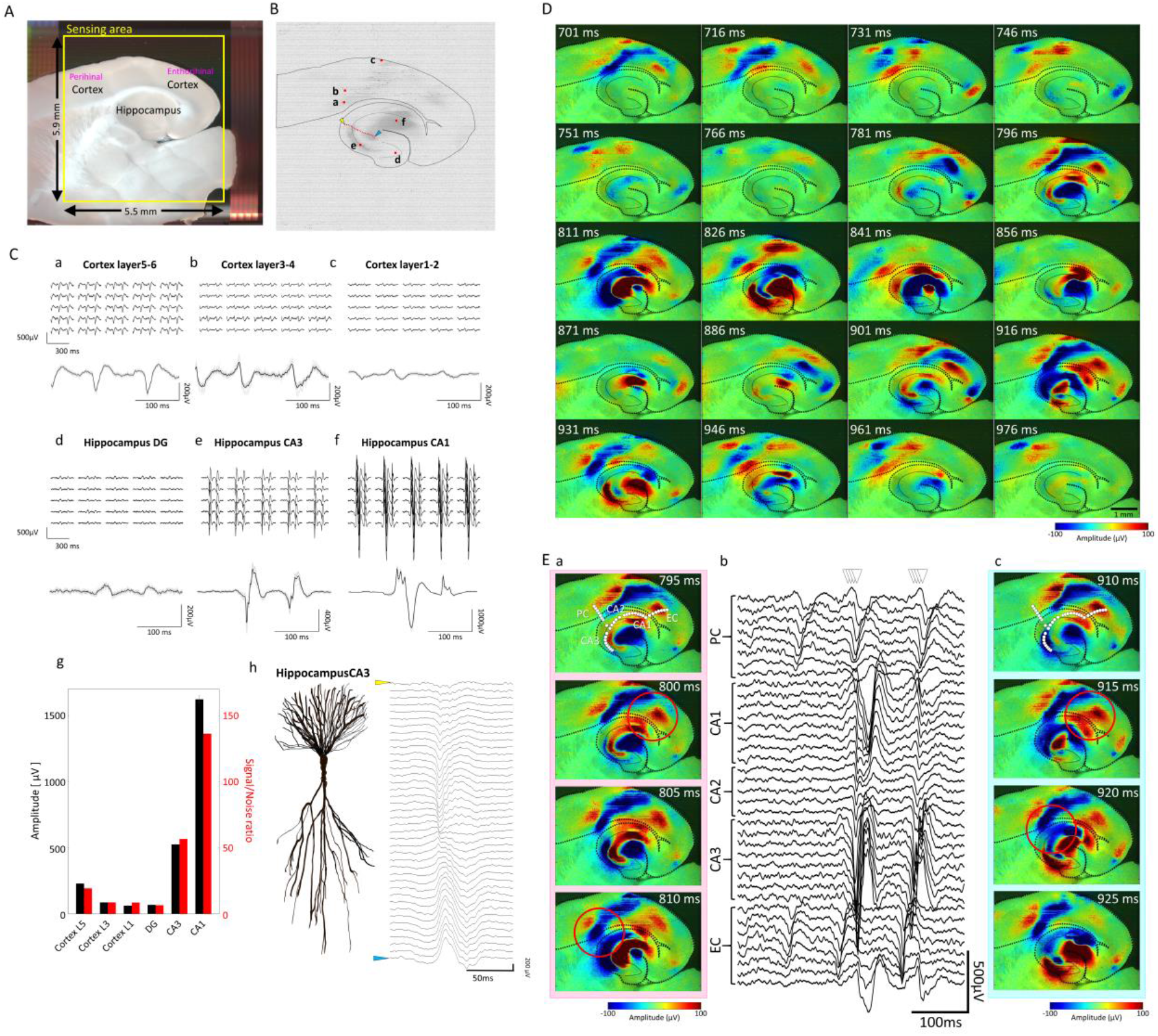
Detailed measurement of spontaneous activity patterns in the hippocampus and cerebral cortex in acute mouse brain slice. (A) Mouse brain slice on HD-CMOS-MEA. The yellow square indicates a sensing area of 5.5 mm × 5.9 mm. (B) Raw waveform in 236,880 electrodes and schematic diagram of cortical and hippocampal areas. (a) Cortical layer 5. (b) Cortical layer 3. (c) Cortical layer 1. (d) Hippocampus DG. (e) CA3. (f) CA1. (C) Raw waveforms in the cortical and hippocampal regions. The waveforms of 25 electrodes in the regions (a) to (f) are indicated by red dots in (B) and the average waveform is a combination of the waveforms of 25 electrodes. (g) Schematic CA3 pyramidal neuron morphology and waveforms acquired at the electrodes indicated by the red line in (B). Waveforms are arranged from yellow to blue arrows in (B). (D) Temporal changes of voltage heatmaps in spontaneous activities in the hippocampus and cortex regions. The propagation pattern for 275 ms is shown as a voltage heat map at 15ms intervals. (E) Synchronized firings between hippocampal and cortical regions. Temporal changes of voltage heatmaps for two synchronous firings first on left (a), second on right (c), for 15 ms at 5 ms interval). The raw waveform of the white dotted line part of the heat map (Hippocampus CA3, CA2, CA1, Perirhinal cortex, Entorhinal cortex regions) was picked up (b). Red circles indicate synchronized firing between hippocampal CA1 and EC region, hippocampal CA3 and PC region.

Surprisingly, due to the high spatiotemporal resolution, the anatomical structures of the hippocampus and cerebral cortex are highlighted from the potential waveforms by positive and negative potential imaging (Figure 2D, Supplementary movie 1). Additionally, we can find the relationship between pyramidal neuronal morphology in CA3 and sink and source in LFP at a certain time. From the radiatum layer to the lacunosum-molecular layer, the potential changed from negative (sink) to positive (source) (Figure 2C–h). Local propagation within layer 5 of the cortex, propagation from the EC to the PC, propagation from the hippocampal DG to CA3 and CA3 to CA1, and input from CA3 to CA1 via Schaffer collaterals were observed along the brain regions. (Figure 2D, Supplementary Movie 1).

We have used low Mg^2+^ 0.1 mM to induce epileptiform activity *in vitro* which resembles Inter-Ictal events (I-IC, Supplementary Movie1, Figure 2D, E) recorded in humans with EEG before or after an epileptic seizure^58, 59^. Figure 2D shows that the cortical region fires independently at around 700 ms, while synchronous firing in the cortical and hippocampal regions was confirmed at about 800 and 900 ms. In both synchronous firings (Figure 2E–b), we observed a phenomenon in which the firing from the EC was linked at the CA1 and CA3 sites. As shown by the circles in Figures 2E(a) and 2E(c), we observed a phenomenon in which the firing from the EC was linked at the CA1 and CA3 sites in both synchronous firings. The projection pathways between EC and hippocampal regions include the cortico-ammonic nucleus pathway with direct input from the EC to the CA1 area, and the trisynaptic pathway via the EC, dentate gyrus, CA3, and CA1 area^60^. Voltage heatmap data was electrical activity consistent with these projection pathways.

### Propagation of gamma waves and sharp-wave ripple (SPW-R) waves in brain slice

The frequency characteristics of LFP have been extensively studied in the hippocampus and cerebral cortex, and are an effective index for evaluating brain function. The gamma frequency has been implicated in cognitive function, and sharp wave-ripple (SPW-R) waves are a marker of epilepsy. They have been reported to have stronger connections with the cortex. Here, we investigated the spatiotemporal distribution of gamma and ripple wave components in the hippocampus and cortex (Figure 3). Gamma wave propagation was observed within the cortex and the hippocampal CA1 and CA3 regions. In the hippocampus, a pattern of propagation from CA3 to CA and CA3 to CA1 was observed (Figure 3A-a). Like gamma waves in the hippocampus, ripple waves propagated between CA3 and CA1 (Figure 3B-a,b). Few ripple waves were observed in the cortex, but interestingly, local oscillations were observed in the cortex on the CA1 side. The result indicates a connection between the hippocampus and the cerebral cortex of ripple waves. Our brain slice measurement by HD-CMOS-MEA was shown investigating the propagation between brain regions according to frequency.

**Figure 3.**
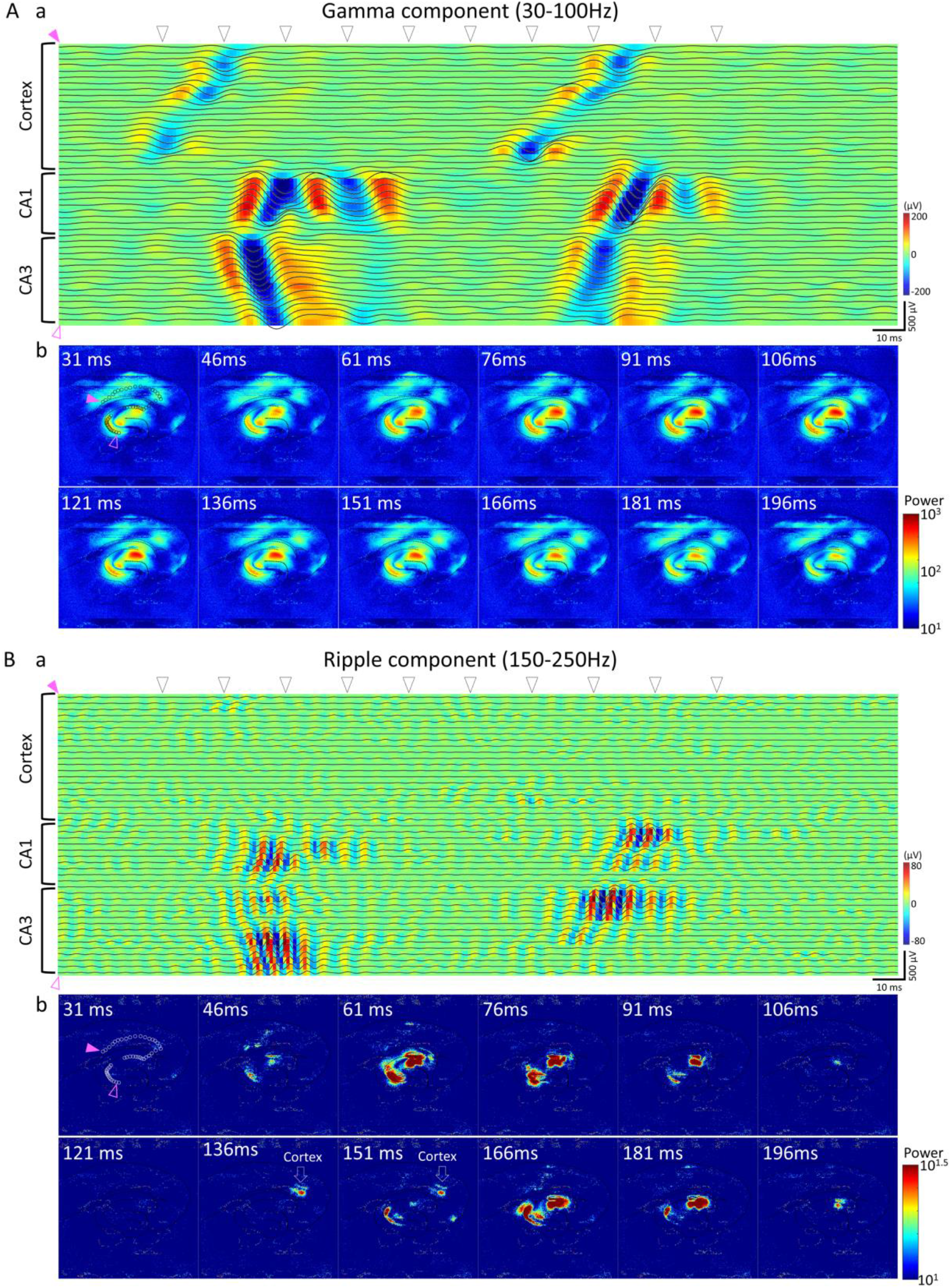
Propagation of Gamma and Ripple waves in hippocampus and cortex. A:(a) Waveforms and voltage heatmaps for 250 ms obtained by band-pass filtering (30-100 Hz) of the signals in the cortical area, CA1, and CA3 areas. (b) Time-lapse of the intensity heatmap (scalogram) of the gamma waveband obtained by wavelet transform. The waveform and voltage heatmaps picked up in (a) the data acquired at the electrode indicated by the black dot in 31 ms image. Pink triangles indicate the starting and ending electrodes. B: (a) Waveforms and voltage heatmaps for 250 ms obtained by band-pass filtering (150–250 Hz) the signals in the cortical area, CA1, and CA3 areas. (b) Time-lapse of scalogram of the ripple waveband. Arrows at 136 ms and 151 ms indicate that ripple components were observed in the cortex area. The triangle arrows in the top of the waveforms and voltage heatmaps (a) indicate times of time-lapse images. The time of the triangle arrows was the same as the triangle arrows in Figure 2E.

### Network analysis and single neuron analysis in drug response of human iPSC-derived cortical neurons

We have evaluated drug responsiveness in hiPSC-derived cortical networks. Figure 4A(a) shows the morphology of hiPSC-derived neurons cultured on 324 electrodes (18 × 18 electrodes). We observed that the soma of a single neuron spanned approximately four electrodes. The locations and firing frequencies of each neuron are shown in Figure 4A(b). The center of the circle represents the centroid position of each neuron, and the size of the circle depicts the firing frequency of each neuron. As a result of spontaneous activity measurement for 1 minute, firing was detected at an average of 285 ± 52 neurons/well (n = 9 wells, Supplementary Movie2). Figure 4A(c) shows the distribution of firing frequency in all neurons (n = 9 wells). The firing frequency ranged from 0.029 to 9.994 Hz, and the mode was 0.9 Hz. These results indicate that HD-CMOS-MEA system can analyze the spontaneous activity of the hiPSC-derived cortical network on a cell-by-cell basis. Figure 4B shows a raster plot for each neuron, a heat map of burst parameters in the network, and a heat map of burst parameters in single neurons, when 4-AP, PTX, and AP5 + CNQX were administered. Black bands in Figure 4B indicate network bursts (NB), and NB was observed in all wells before drug administration. 4-AP administration increased the total spikes (TS) ant total spikes per single neuron [TS(s)], NBs, and burst in a single neuron (B), and decreased the CV of spikes in an NB and Max Frequency in a B (MF) (Figure 4B-a). The administration of picrotoxin (PTX), a GABA-A receptor blocker, increased the duration in NB and B, the spikes in B at 1 µM (Figure 4B-b). It was confirmed that GABA-A receptors are functionally expressed in hiPSC-derived neural networks. These results also indicate that both the network and single neuron analyses can detect the response of the seizurogenic compounds.

**Figure 4.**
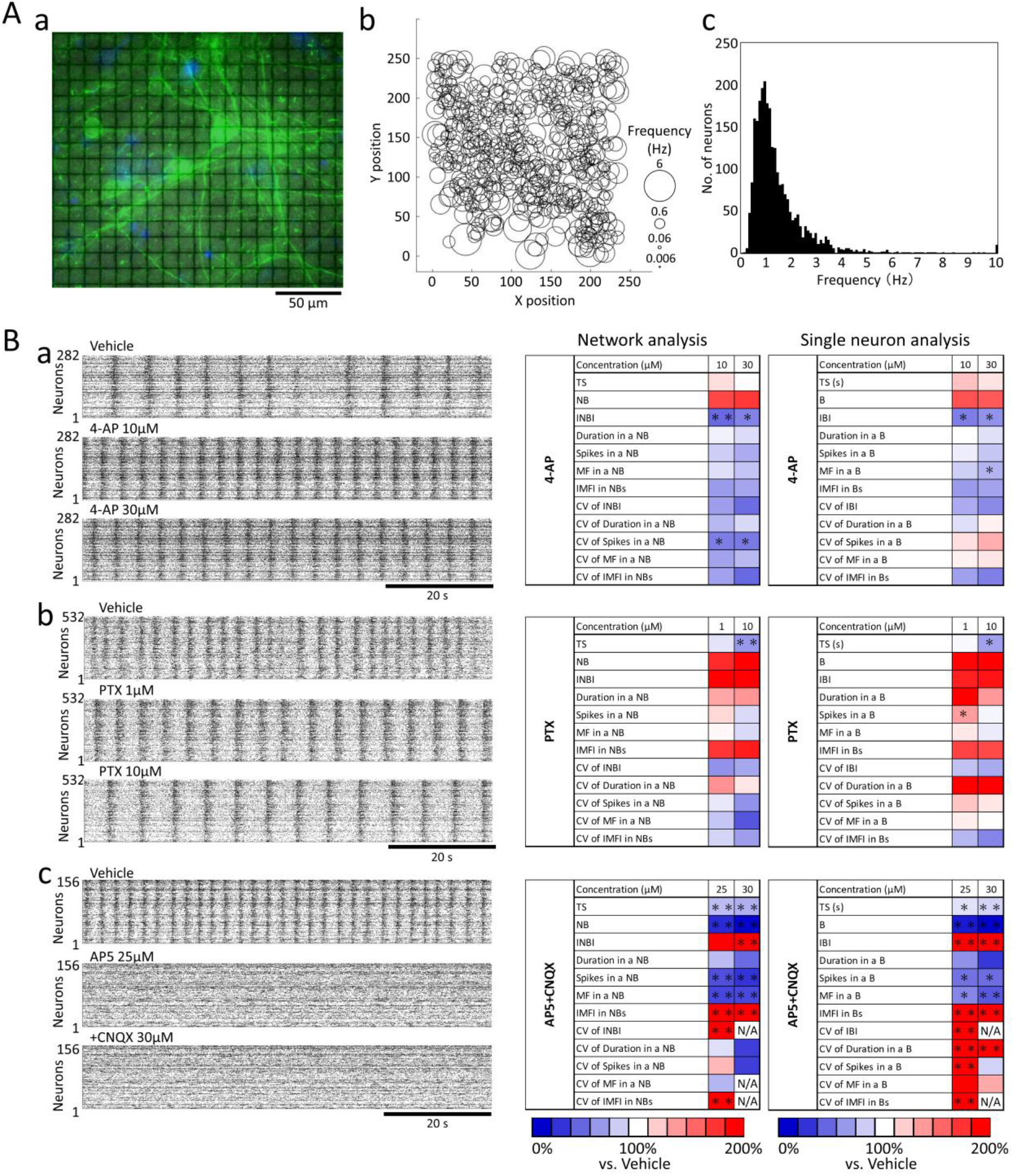
Network analysis and single neuron analysis to compounds responses in hiPSC-derived cortical neurons. (A) Spontaneous activity frequency of human iPS cell-derived cortical neurons per neuron. Spontaneous activity measurement and pharmacological tests were carried out after 42 days of culture. (a) Immunostaining image of human iPS cell-derived neurons cultured on CMOS-MEA (18 × 18 = 324 electrodes) at six weeks of culture. Green: β-tubuliine III, Blue: Hoechest33258. (b) Spontaneous activity frequency per neuron detected by 235 × 252 = 59,220 electrodes. The size of the circle indicates the spontaneous firing frequency of 532 neurons. (c) Histogram of spontaneous activity frequency per neuron (n = 9 CMOS-MEAs, n = 2569 neurons). (B) Representative raster plots and heatmaps of network analysis and single neuron analysis for drug administration. (a) Vehicle, 4-AP 10 µM, 30 µM. (b) Vehicle, PTX 1 µM, 10 µM. (c) Vehicle, AP-5 25 µM AP-5+CNQX 30 µM. Heat map shows % versus vehicle = 100% (n = 3, *: p ≦ 0.05, **: p ≦ 0.01 vs. vehicle, Dunnett’s test). N/A: unable to calculate due to loss of burst. Parameters in network analysis [TS: Total spikes, NB: Network burst, INBI: Inter NB Interval, Duration in a NB, Spikes in a NB, MF in a NB: Maximum frequency in a NB, IMFI in NBs: Inter MF Interval in NBs, CV of INBI: Coefficient of variance (CV) of Inter NB Interval, CV of Duration in a NB, CV of spikes in a NB, CV of MF in a NB, CV of IMFI in NBs]. Parameters in single neuron analysis [TS(S): Total spikes per single neuron, B: Burst, IBI: Inter burst Interval, Duration in a B, Spikes in a B, MF in a B: Maximum frequency in a B, IMFI in Bs: Inter MF Interval in Bs, CV of IBI: Coefficient of variance (CV) of Inter B Interval, CV of Duration in a B, CV of spikes in a B, CV of MF in a B, CV of IMFI in Bs.]

AP5 (25 µM), an antagonist of NMDA-type glutamate receptors, decreased the 5 parameters (TS and TS(S), NB and B, duration in a NB and B, Spikes in a NB and B, MF in a NB and B), and increased 3 parameters [Inter MF Interval (IMFI) in NB and B, CV of Inter NB and B Interval, CV of IMFI in NB and B]. CV of duration in a B and CV of spikes in a B changed only in single neuron analysis. Additional administration of CNQX 30 µM, an AMPA receptor inhibitor, also decreased the 5 parameters and increased 2 parameters (Inter NB and B interval, IMFI in NB and B). CV of INBI and IBI, CV of MF in a NB, CV of IMFI in a NM and B were not calculated because CNQX reduced or eliminated NB and B (Figure 4B-c). From these results, it was confirmed that the function of both NMDA and AMPA-type glutamate receptors in cultured hiPSC-derived cortical neurons.

Interestingly, the coefficient variance (CV) of duration in an NB and B, CV of spikes in an NB and B, and CV of MF in an NB and B were different between network analysis and single neuron analysis of four compounds. These results indicate that adding single neuron analysis parameters increases the amount of information and provides new information in evaluating the drug responsiveness of neural networks.

### Evaluation of synaptic strength by compound administration

To investigate changes in synaptic strength due to the compound administration, firing time-series data of each neuron was extracted, and the number of spikes fired within 100 ms for each spike of each neuron (synchronized spike) was counted for all combinations of neurons. Next, 100 surrogate time-series data were created by randomly rearranging the inter spike intervals (ISI) calculated from the time-series data of each neuron, and synchronized spikes were counted. Figure 5A-a shows synchronized spikes (red bar) fired within 100 ms with neuron1 as a reference for real and surrogate time-series data. Z score was calculated from the real synchronized spikes and 100 surrogate synchronized spikes. Figure 5A-b shows histograms of synchronized spikes from the combination of Neuorn1 and Neuron2, and Neuorn1 and Neuron3. The black bar represents 100 surrogate data, and the red represents the real data. The Z score for neuron1 versus neuron2 was 3.52, and for neurons1 versus neuron3 was −3.80. When the Z score of the real synchronized spike exceeded 3, the combination of neurons was recognized as an excitable connection. Conversely, when the Z score was less than −3, it was recognized as an inhibitory connection. The color map on the left in Figure 5B-a,b,c shows the Z score matrix between each neuron, and the middle shows the Z score histogram for all connections. The red area of the histogram indicates an excitable connection, and the blue area indicates an inhibitory connection. Before compound administration, the number of combinations with a Z score of ≧3 was 33.5% ± 4.3% (n = 9 wells), which indicates regular excitatory connections were formed. For 4-AP, the percentage of excitability binding tended to decrease, −1.08% ± 4.02.% at 10 µM and −4.25% ± 5.30% at 30 µM, and the inhibitory connections changed slightly, 0.176% ± 0.20% at 10 µM., −0.66% ± 0.39% at 30 µM (Figure 5B-a). On the other hand, PTX increased the percentage of excitability connections, 8.44% ± 1.80% (p < 0.01) at 1 µM and 5.38 ± 0.47% (p < 0.05) at 10 µM (Figure 5B-b). The inhibitory connections changed slightly, −0.63% ± 0.35% at 1 µM and −1.17% ± 0.76% at 10 µM (Figure 5B-b). These results indicate that PTX enhances excitatory coupling and shifts to a regular firing pattern compared to 4-AP. Administration of 25 µM AP5 decreased the percentage of excitability connections to −39.04% ± 7.68%, and the subsequent addition of 30 µM CNQX did not change the percentage, which was −39.06% ± 7.62% (Figure 5Cc). Inhibitory connection was unchanged, −0.08% ± 0.27% at AP5 25 µM and 0.07% ± 0.27% at +CNQX 30 µM. This indicates that excitatory synaptic connections were blocked, which results in random firing.

**Figure 5.**
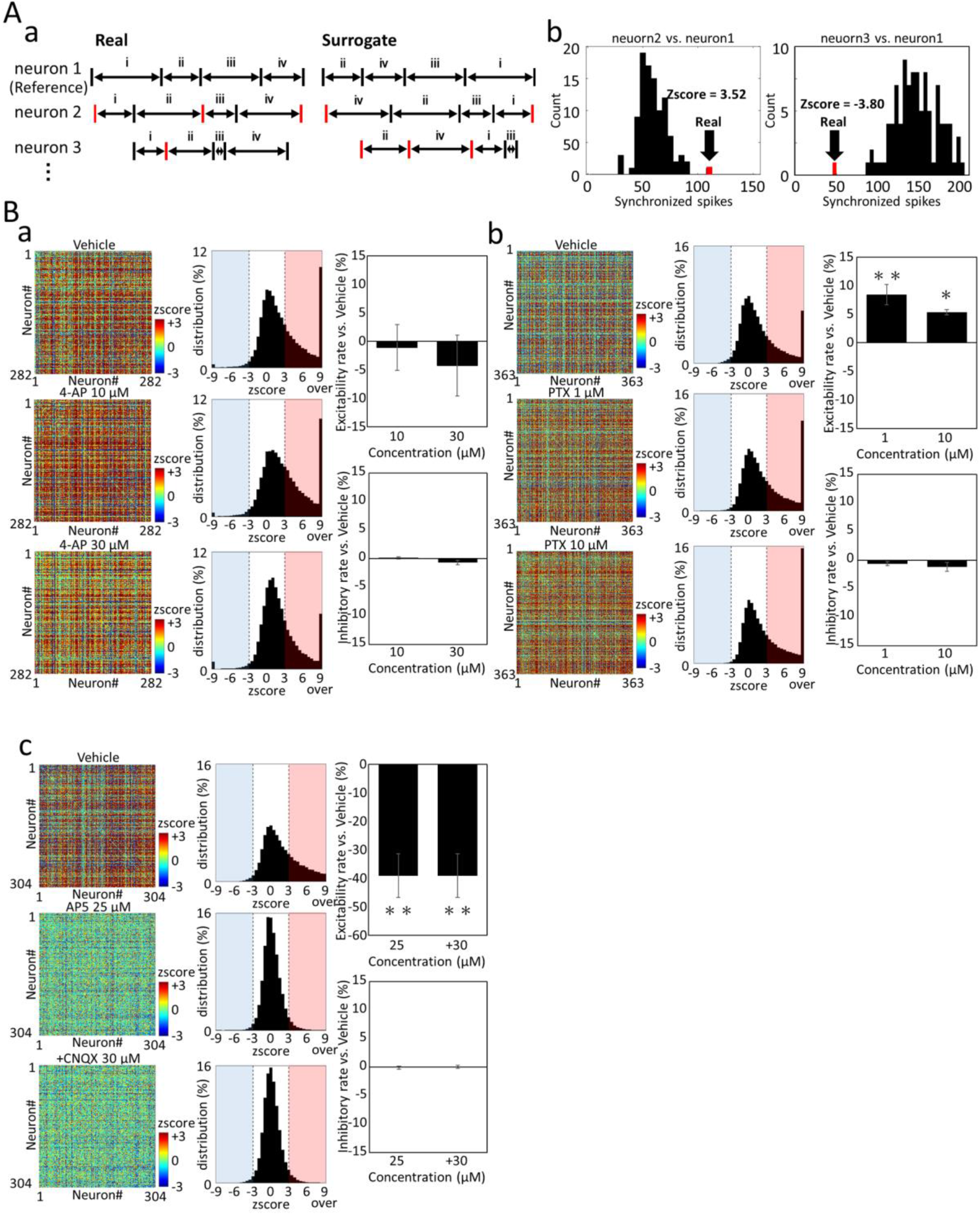
Changes in the synaptic strength due to compound administration in hiPSC-derived cortical neurons. (A) Z score calculation to quantify connection strength between neurons. (a) Raster plot of three representative neurons. Left: real data, right: random surrogate data. Red raster plots show spikes that fire within 100 ms of neuron1 firing. (b) Synchronized spike histogram calculated from spike trains of neuron2 and neuron1 (left) and synchronized spike histogram calculated from spike trains of neuron3 and neuron1 (right). Black indicates random surrogate data, red indicates a synchronized spike of real data. (B) Changes in Z score due to compounds administration. (a) vehicle (Top), 4-AP 10 µM (middle), 4-AP 100 µM (Low). (b) vehicle (Top), PTX 1 µM (middle), PTX 10 µM (Low). (c) Vehicle(Top), AP-5 25 µM (middle), AP-5 25 µM+CNQX 30 µM (Low). Left: Matrix of Z scores between each neuron before and after compound administration. The color scale is −3 ≤ zscore ≤ 3. Middle: Z score histogram of all connections. The red area (Z score ≥ 3) indicates an excitable connection, and the blue area (Z score ≤ 3) indicates inhibitory connection. Right: Excitability connections percentage change (top, Ave. ±S.E., n = 3) and inhibitory connections percentage change (low, Ave. ±S.E., n = 3) due to compounds administration (* : p ≦ 0.05, ** : p ≦ 0.01 vs. vehicle, Dunnett’s test).

Among the difference in reactivity between 4-AP and PTX, we calculated the changes in the firing frequency of one neuron and the proportion of connections (with a Z score of 3 or more) between one neuron and other neurons (excitability connection rate: ECR, ECR = number of Z score ≧3/number of neurons-1*100%) (Figure 6). Figure 6A-a shows the distribution of the firing frequency of single cells before and after administration of 4-AP (n = 193 neurons, maximum firing frequency: 6.47 Hz) and changes in ECR. The size of the dots indicates the firing frequency, and the color indicates the increase/decrease of the ECR for the vehicle (100%). At 10 µM of 4-AP, the percentage of neurons with increased ECR was 22.8% and the percentage of decrease was 75.6%, and at 30 µM, the percentage of neurons with increased ECR was 50.8% and the percentage of decrease was 47.7%. The rate of unchanged ECR was 1.6% for both 10 µM and 30 µM. At low concentrations, the ECR of about 80% of the cells decreased; at high concentrations, the rate of increase and decrease was approximately similar. Looking at the size of the circles in Figure 6A-a, there was no significant finding in firing frequency of each neuron after 4-AP administration. Figure 6A-b shows distribution maps of connection rates after 4-AP administration. The vertical axis represents the ECR after administration, the horizontal axis represents the ECR during the vehicle, and the dotted line represents the line with no change. 4-AP 10 µM showed a tendency to reduce neurons with a vehicle ECR of 50% or more, and 4-AP 30 µM showed a tendency for increasing neurons and decreasing neurons sparsely distributed regardless of the vehicle ECR. Figure 6B-b shows the distribution of single-cell firing frequency (n = 363 neurons, maximum firing frequency: 7.24 Hz) and changes in ECR before and after PTX administration. At 1 µM PTX, the percentage of neurons with increased ECR was 76.9%, the percentage of decrease was 22.3%, and the percentage of neurons without change was 0.8%. At 10 µM, the percentage of neurons with increased ECR was 76.9%, the percentage of decreased was 22.0%, and the percentage of neurons without change was 1.1%. PTX increased the ECR about 80% of the neurons from low concentrations, and the tendency was almost the same even at high concentrations. Additionally, changes in the firing frequency of a single neuron were observed to change compared to 4-AP (Figure 6B-a). Figure 6B-b shows the distribution map of the connection rate after PTX administration. At both PTX 1 µM and PTX 10 µM, many neurons showed an increase in ECR, and the range of increase in neurons with a vehicle ECR of 20% or less was extensive. PTX 10 µM tended to have a more extensive ECR fluctuation range than 1 µM. Compared to 4-AP, PTX increased the number of neurons with increased ECR with change in firing frequency of single neurons.

**Figure 6.**
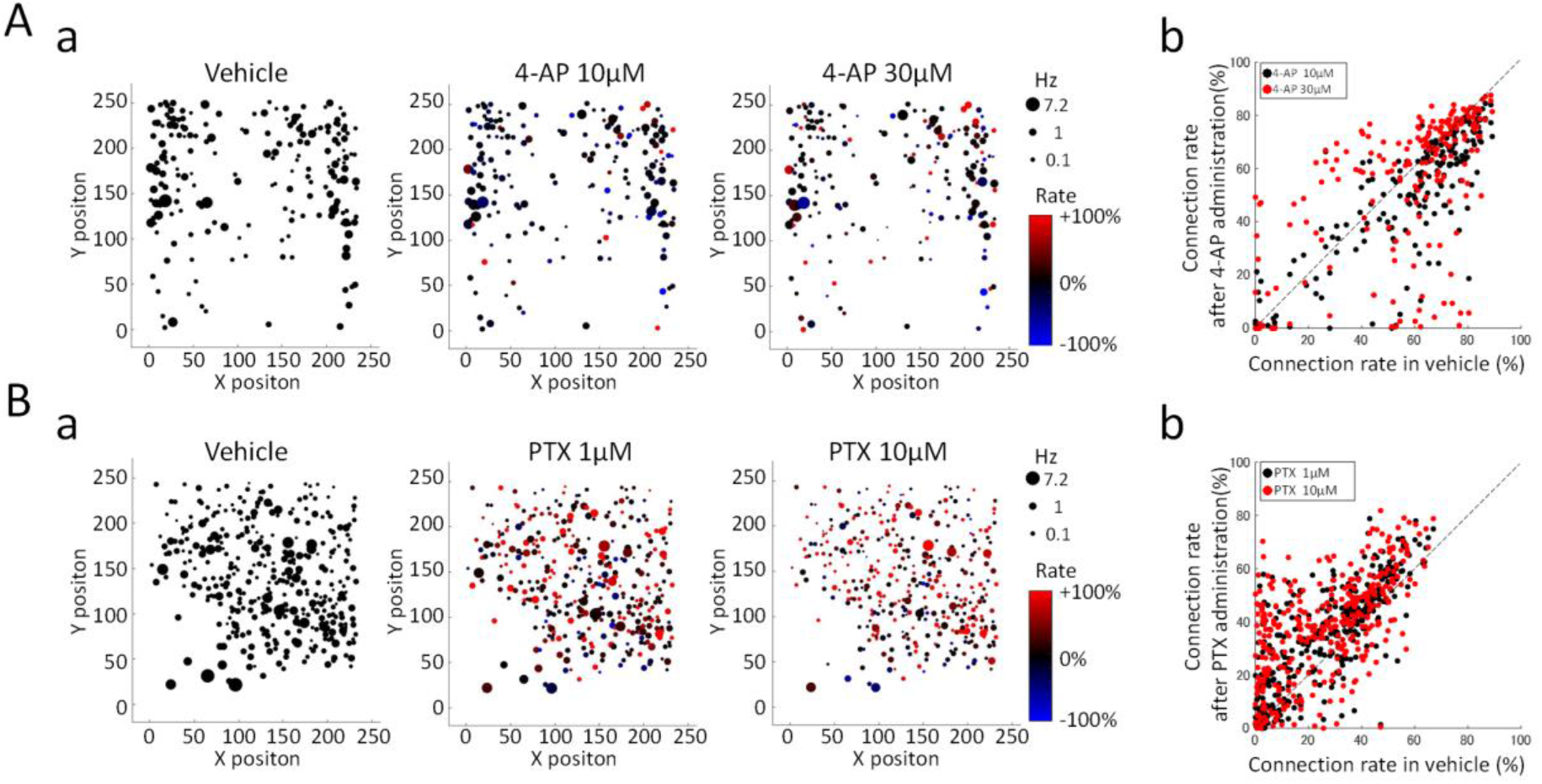
Changes in firing frequency of single neuron and excitability connection rate (ECR) after 4-AP and PTX administration. There were positions of neurons where firing was detected on the 59,220 electrodes (235 × 252 electrodes) and firing frequencies in single neurons. The firing frequency is indicated by the size of the circle. The size of the circle indicates the firing frequency. The ECR value of each neuron in vehicle is defined as 100%, and the ECR increase/decrease due to drug administration is shown in color. (A-a) Vehicle, 4-AP 10 µM, 30 µM. (B-a) Vehicle, PTX 1 µM, 10 µM. (A-b) 4-AP 10 µM, 30 µM. (B-b) Vehicle, PTX 1 µM, 10 µM.

These results showed that the analysis of the intercellular Z score calculated from the firing timing of single neuron is an analysis method for changes in the synaptic strength.

### Spontaneous activity pattern of DRG neurons on a cell-by-cell basis

We have evaluated the detection of cell-by-cell spontaneous activity patterns of DRG neurons and their responsiveness to pain-related compounds. Figure 7A-a shows DRG neurons cultured on HD-CMOS-MEA at 6 WIV (weeks in vitro). A peculiar appearance of DRG neurons with different soma sizes neurites was observed on HD-CMOS-MEA. The soma of about 100 μm in the major axis direction was located on 29 electrodes (Figure 7A-b). The spontaneous activity of each neuron was measured for 60 seconds, and the frequency of each neuron with spontaneous activity was detected (n = 993 neurons, n = 7 wells). Figure 7A-c shows the soma position and firing frequency detected on one HD-CMOS-MEA in the size of a circle. Identification of soma location included neurons that responded to capsaicin (Figure 8). Therefore, neurons that were not spontaneously active were identified at the time of spontaneous activity measurement. Figure 7A-d shows the distribution of spontaneous activity frequencies of all 993 neurons. The number of nonspontaneous neurons was the highest at 35.65% (n = 354 neurons), and the number of neurons that were frequently spontaneously active above 10 Hz was also observed at 4.63% (n = 46 neurons). Unlike the central nervous system, DRG neurons have no synchronous activity and are spontaneous activities without synaptic transmission^41^. Figure 7B shows a 1-minute firing histogram of typical neurons firing in each frequency band. The proportion of nonspontaneous neurons is highest at 35.65% (n = 354 neurons), 0 < frequency (F) < 0.1 Hz neurons 23.97% (n = 238 neurons), 0.1 ≤ F < 0.5 Hz neurons 18.63% (n = 185 neurons), 0.5 ≤ F < 1 Hz neurons were 6.34% (n = 63 neurons), 1 Hz ≤ F < 2 Hz neurons were 4.53% (n = 452 neurons), 2 Hz ≤ F < 5 Hz neurons were 4.03% (n = 40 neurons), and 5 Hz ≤ F < 10 Hz neurons were 2.22% (n = 22 neurons). Frequently spontaneously active neurons above 10 Hz were present at a rate of 4.63% (n = 46 neurons). The maximum firing frequency was 113.1 Hz. We found that there are DRG neurons which exhibit high-frequency firing above 100 Hz. Based on these results, it was found that DRG neurons have spontaneous activity with different firing frequencies.

**Figure 7.**
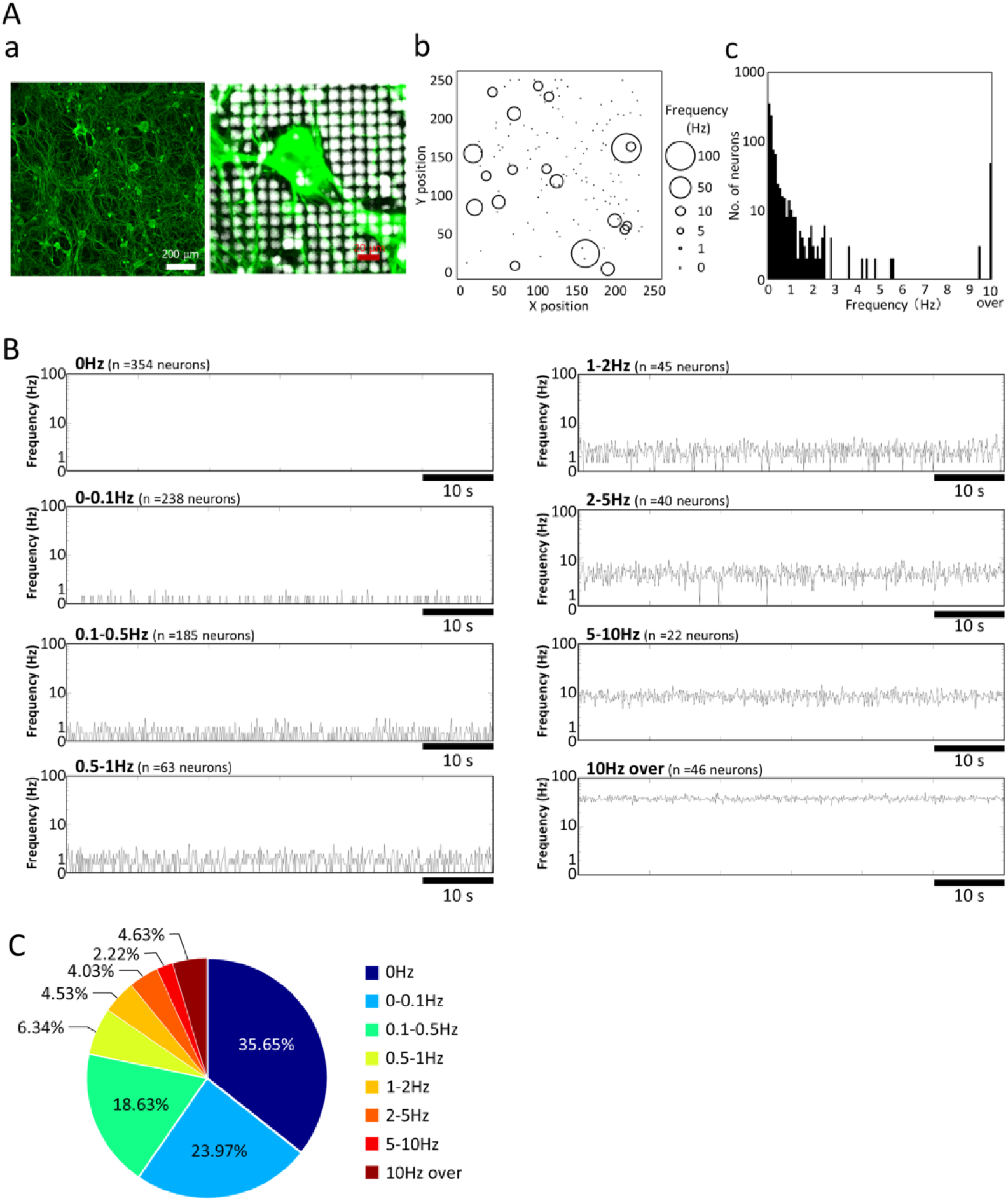
Spontaneous activity frequency distribution in cultured DRG neurons. (A) Immunostaining image of DRG neurons (β-tubuline III) at 6 WIV (weeks in vitro) cultured on HD-CMOS-MEA (a). Enlarged view near the soma (b). DRG neurons detected on the 235 (X-axis) x 252 (Y-axis) = 59,220 electrodes and spontaneous activity frequency. The size of the circle indicates the frequency of spontaneous activity from 0 Hz to 100 Hz (c). Histogram of spontaneous activity frequencies of DRG neurons (bin = 100 ms, n = 993 neurons, n = 7 HD-CMOS-MEAs) (d). (B) The spontaneous activity of single DRG neurons is a typical firing pattern per minute in each frequency band. (C) Percentage of DRG neurons in each spontaneous firing frequency band.

**Figure 8.**
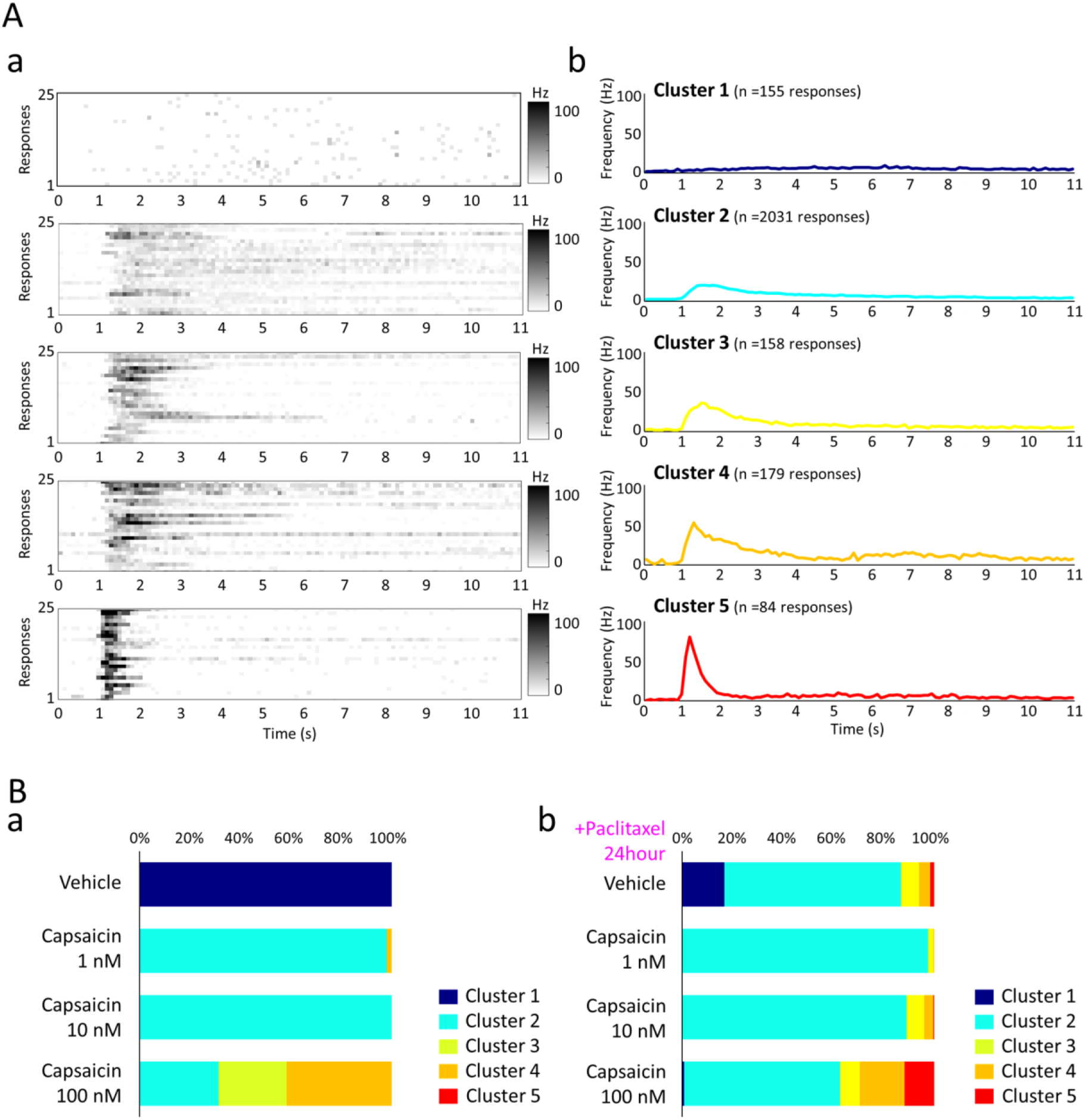
Responses to capsaicin in the presence and absence of paclitaxel. (A) Clustering of evoked responses in capsaicin administration. Induced response of 25 typical neurons in each of the 5 clusters (bin = 100 ms). The firing frequency of 0 to 100 Hz is shown in gray scale (a). Time course of average firing frequency for each of the 5 clusters (b). (B) Cluster ratio of evoked response to capsaicin 1, 10, 100 nM administration (a). Cluster ratio of evoked response to capsaicin 1, 10, 100 nM administration in the presence of paclitaxel 10 nM for 24 hours (b).

### Response to capsaicin in DRG neurons and enhancing effect by paclitaxel

Next, the evoked response by the administration of capsaicin, an agonist of the TRPV1 channel, was examined. Figure 8A shows the results of clustering the evoked response patterns after capsaicin administration (n = 2607 responses, n = 653 neurons, n = 5 CMOS-MEAs). In each cluster, the evoked response of 25 typical neurons is shown (Fig. 8A-a), and the histogram of the average evoked response is shown in Fig. 8A-b. Cluster1 is a sample group that does not show a rapid increase in the number of firings after administration. Cluster2 had a peak frequency of 20.7 Hz after 1.5 seconds and an average frequency of 7.7 Hz for 10 seconds. Cluster3 had a peak frequency of 35.8 Hz after 1.5 seconds and an average frequency of 9.4 Hz for 10 seconds. Cluster4 had a peak frequency of 52.8 Hz after 1.3 seconds and an average frequency of 13.2 Hz for 10 seconds. Cluster 5 showed a strong peak frequency of 80.2 Hz after 1.2 seconds, after which the firings decreased sharply, with an average frequency of 8.2 Hz for 10 seconds. These results indicate that each DRG neuron has a different evoked response to capsaicin. Figure 8B-a shows the concentration-dependent response of capsaicin. With Capsaicin 1 nM administration, Cluster 2 was 97.87%, and with 10 nM it was 100%, showing the same response. On the other hand, when 100 nM was administered, highly reactive cluster 3 accounted for 27.08% and cluster 4 accounted for 41.67%. Figure 8B-b shows the capsaicin response results after exposure to the anti-cancer drug paclitaxel 3 nM for 24 hours. In the vehicle, cluster 2 was 70.10%, cluster 3 was 7.31%, cluster 4 was 4.32%, and cluster 5 was 1.66%. In capsaicin 1 nM, cluster 2 was 97.36%, cluster 3 was 1.98%, cluster 4 was 0.50%, and cluster 5 was 0.00%. In capsaicin 10 nM, cluster 2 was 88.76%, cluster 3 was 6.94%, cluster 4 was 3.64%, and cluster 5 was 0.50%. With capsaicin 100 nM, cluster 2 was 61.86%, cluster 3 was 7.79%, cluster 4 was 17.74%, and cluster 5, which showed the strongest response at 11.77%, and activation of the TRPV1 channel by paclitaxel administration. The analysis results of the evoked response on a cell-by-cell basis suggested that not all cells showed a uniformly strong response, and that the ratio of evoked response characteristics was related to pain intensity. It was also shown that the effect of anticancer drugs could be evaluated from the evoked response pattern of each neuron.

### Axon conduction measurements

Axon conduction in DRG neurons was measured. Supplementary Movie 3 is an axon conduction movie of action potential emitted from soma of a single DRG neuron at 35 kHz sampling. In the soma, a strong downward voltage with an amplitude of 1,038 µV was observed (Figure 9A-a), and then a couple of positive and negative voltages was observed to conduct axons (Figure 9A-b–f). It shows the sink and source of Na^+^ ions in axons. A heat map presents the time-delayed waveform and the position of the axon having the waveform was observed (Figure 9A). Figure 9B shows the axon conduction pathway with a time-delayed heat map, showing the data at the 4 and 6 WIV. It can be seen that the action potential of the acquired axon is long and widely known at the 6 WIV. Axon conduction in a single DRG neuron could be obtained with more than 500 electrodes. The axons extending from the perikaryon were defined as origin, middle, and terminal in the order of proximity to the perikaryon, and the conduction velocity was calculated from the path, shown on the right in Figure 9B and the delay time at the electrode indicated by the arrow. The axon conduction velocities were 1.39 m/s (origin), 1.20 m/s (middle), 0.77 m/s (terminal) at the 4 WIV, and 1.25 m/s (origin), 1.20 m/s (middle), and 1.01 m/s (terminal) at the 6 WIV, respectively (Figure 9A-c). It was found that the origin was the fastest in both the 4th and 6th weeks of culture, and the conduction velocity decreased toward the end. These results indicate a decrease in conduction velocity in the unmyelinated axons.

**Figure 9.**
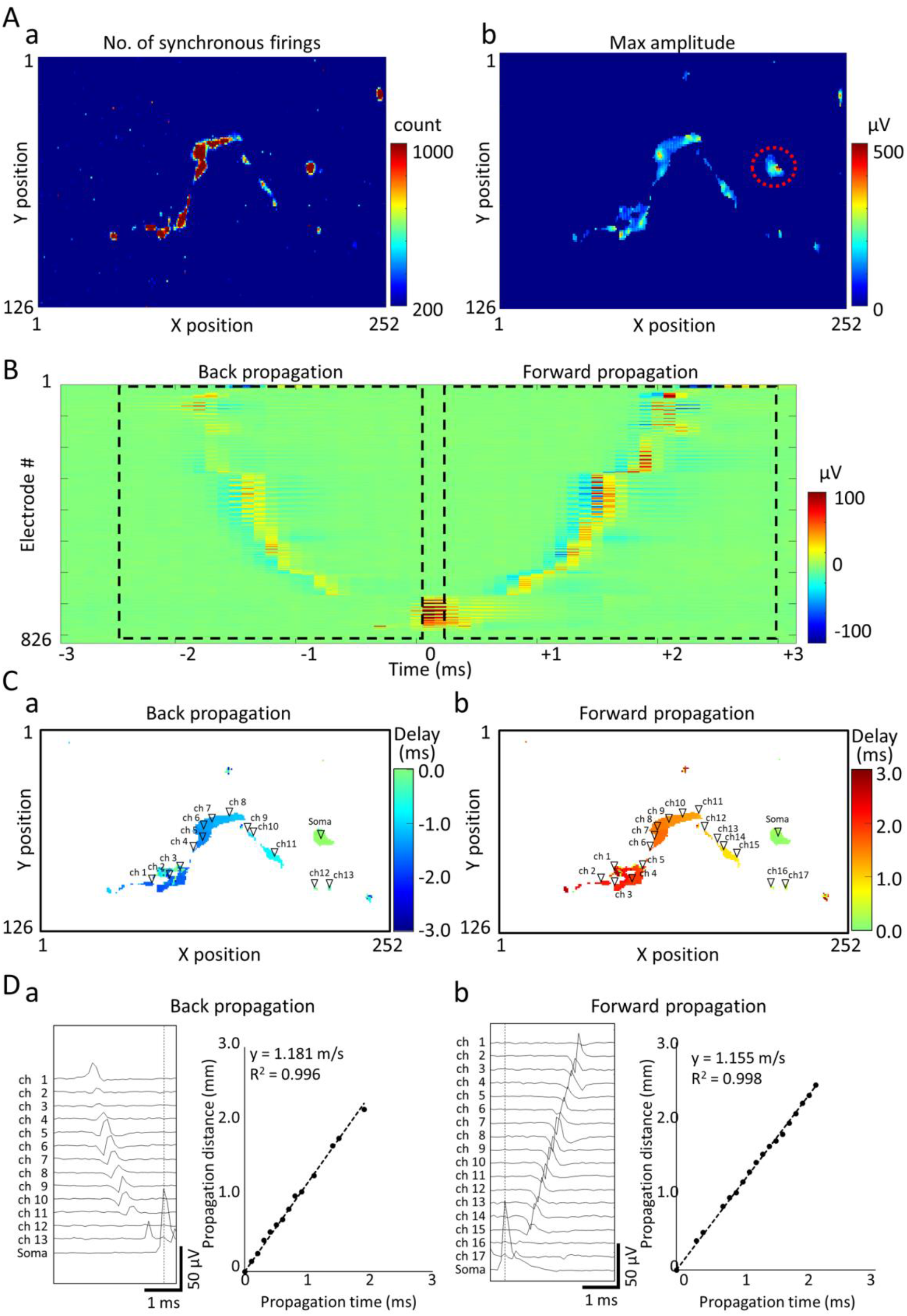
Back propagation and forward propagation in single axonal conduction. (A) Identification of soma electrode locations, (a) Number of synchronous firings map, (b) Maximum amplitude map of electrodes with more than 200 synchronous firings detected The dotted line indicates the soma electrode area specified from the voltage value and the shape of the electrode distribution. (B) Average potential heat map for 3 ms before and after soma firing. (C) Conduction path and delay time map, (a) Heat map of back conduction 3 ms before soma firing, (b) Heat map of forward conduction 3 ms after soma firing. (D) Electrode voltage waveform used for conduction velocity calculation (left). The dotted line indicates the firing time of the soma. Plot of conduction time and cumulative conduction distance (right). The dotted line is the approximate straight line calculated by the least squares method. (a) Back conduction, (b) Forward conduction.

### Back propagation and forward propagation in axon

To identify the conduction path of neurons, the maximum amplitude of each electrode was detected every 0.05 seconds. Then, synchronous firings were defined as maximum amplitudes with a time delay of less than 1 ms for 4 or more among the 8 electrodes. We counted the synchronous firings of all electrodes from 76.8 seconds of data, 826 electrodes with more than 200 synchronous firings were derived for analysis (Figure 9A-a). Next, we created a maximum voltage amplitude map for 76.8 seconds of 826 electrodes and determined the soma location from the voltage values and the circular shape of the electrode distribution (Figure 9A-b). 100 electrodes detected soma firing, and the maximum amplitude was 521.2 µV. Spike detection of the electrodes (100 electrodes) where the soma is located was performed using a voltage threshold of 200 µV, and 1,167 firings were detected. We found a neuron with high-frequency spontaneous firing at 15.2 Hz. By using the firing time of the soma as a reference, voltage values were extracted 3 ms before and after firing, and the average voltage map of 1,167 times was calculated (Figure 9B). Forward propagation was observed for 3 ms after soma firing. Propagation was also observed for about 2 ms before firing the soma. The forward propagation and back propagation had the same route (Supplementary movie 4). Figure 9C shows the conduction pathways around the time of soma firing as a time heat map was shown that the paths of back propagation and forward propagation are the same. Starting from the position of the center of gravity of the soma, the farthest electrode at which firing was obtained every 0.1 ms was selected, and the cumulative linear distance between the electrodes was calculated as the conduction distance. The figure shows the conduction waveforms and conduction velocities of backpropagation and forward propagation. In the waveform, the negative potential (sink) appeared first, followed by the positive potential (source) (Figure 9B, D). The back propagation distance was 2.18 mm, the conduction time was 1.9 ms, and the conduction velocity was 1.181 m/s (R^2^ = 0.996). Forward propagation distance was 2.44 mm, conduction time was 2.1 ms, and conduction velocity was 1.155 m/s (R^2^ = 0.998). Back propagation and forward propagation had almost the same speed.

### Change of axonal conduction velocity to anticancer drug vincristine

Next, we examined how the axonal forward propagation velocity changes with the administration of an anticancer drug, vincristine. Figure 10 (a) shows typical forward propagation patterns before and after vincristine 3 nM administration. The number of electrodes detect forward propagation was 719 electrodes before, 436 electrodes 2 hours after administration, and 504 electrodes 24 hours after administration. A graph of conduction distance versus conduction time for this axon is presented in Figure 10 (b). The conduction velocity was 1.16 m/s (2.44 mm/2.1 ms) before administration, 1.37 m/s (2.05 mm / 1.5 ms) after 2 hours, and 0.85 m/s (1.70 mm / 2.0 ms) after 24 hours. Figure 10(c) shows changes in axonal conduction velocities acquired with separate CMOS-MEAs (n = 4 axons, n = 4 wells). Conduction velocity increased by 121.0% ± 7.1% (p = 0.0234) 2 hours after administration of vincristine and by 109.7% ± 4.4% (p = 0.3066) 24 hours after administration of vincristine. Two hours after the administration of vincristine 3 nM, axonal conduction velocity increased.

**Figure 10.**
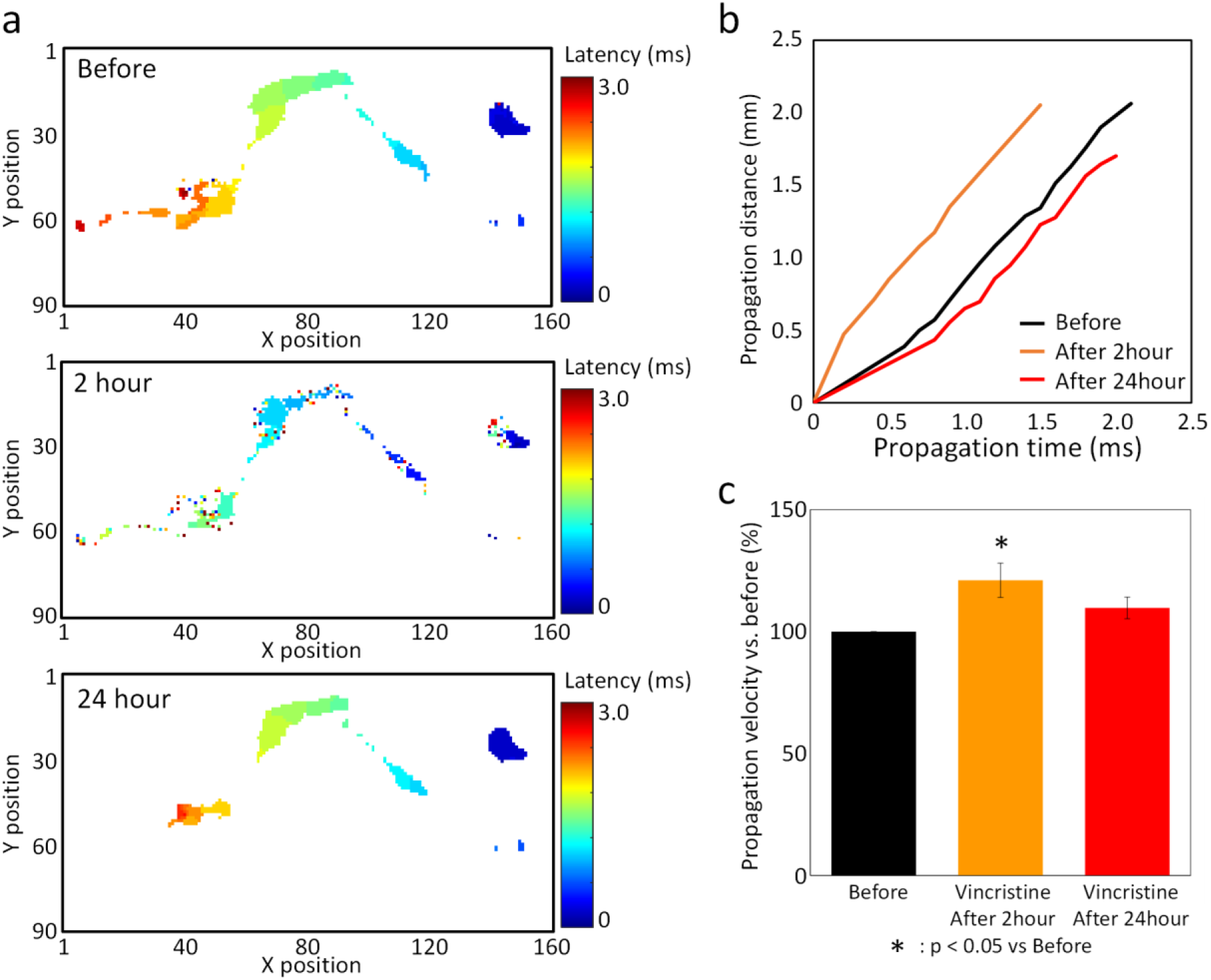
Change of axonal conduction velocity to anticancer drug vincristine. (A) Heat map of forward propagation 3 ms after soma firing before and after vincristine administration. The upper row is before administration, the middle row is 2 hours after administration, and the lower row is 24 hours after administration. (B) Plot of conduction time and cumulative conduction distance before and after vincristine administration. (C) Change in the conduction velocity by vincristine. The parameters were depicted as the average % change of before (before set to 100%) ± SEM from n = 4 wells. Data were analyzed using one-way ANOVA followed by post hoc Dunnett’s test (*p < 0.05 vs. before).

### Detection of spontaneous firing patterns and drug responses in patient-derived cerebral organoids

The electrical activity of human cerebral organoids prepared from iPS cells derived from Rett syndrome patients was measured using a HD-CMOS-MEA. The spontaneous activities of organoids cultured for 4 months were measured before and after administration of 4-AP 100 µM. Figure 11A shows a heat map of the number of firings detected in the spontaneous firings of organoids for 153.6 seconds. The area where organoids adhered to the electrodes occupied 14,612 electrodes. Before 4-AP administration, firing was detected from 4,605 electrodes out of 14,612 electrodes, and activity could be acquired from 31.5% of the electrode area. After 4-AP administration, active electrodes increased to 7,260 electrodes (49.7%) (Supplementary movie 5). Figure 11B shows a raster plot of 14,612 electrodes and a histogram of the number of firings (bin = 100 ms). Stripped black lines in raster plots represent network burst firings. Figure 11C shows a 3D raster plot of the organoid activity for 3 seconds indicated by the red line in Figure 11B. As a result of detecting the number of firing electrodes per 5 ms, the maximum number of electrodes before 4-AP administration was 283 electrodes (average 31.8 electrodes). At the same time, the maximum number after 4-AP administration was 339 electrodes (average 45.5 electrodes). An increase in the maximum number of electrodes indicates an enhancement of network-wide synchrony within the burst firings. An increase in the average number of electrodes suggests an enhancement of local synchronized firing at the single-cell level. (Figure 11C). Figure 11D shows the total number of firings, the number of active electrodes, and the average firing frequency per electrode before and after 4-AP administration. 4-AP increased the total number of firings by 307.0% (Vehicle: 286,042 spikes; 4-AP: 878,366 spikes) and the number of active electrodes by 157.6% (Vehicle: 4,605 electrodes; 4-AP: 7,260 electrodes). The average firing frequency per electrode increased by 194.8% (Vehicle: 0.40 Hz, 4-AP: 0.79 Hz). In summary, we found that it is possible to acquire a wide range of spontaneous activities of human brain organoids, detect changes in detailed network burst patterns, and detect changes due to drug administration.

**Figure 11.**
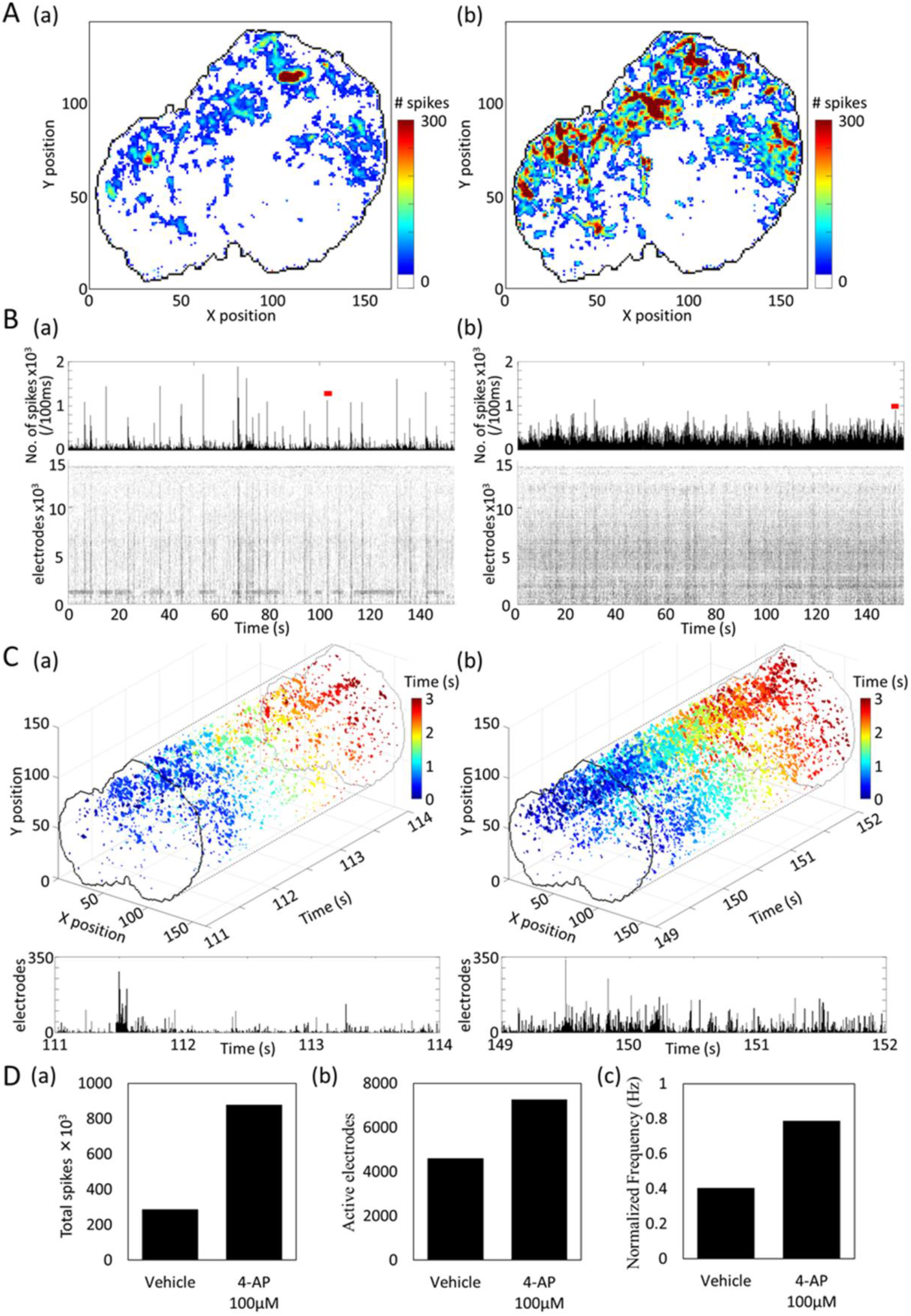
Spontaneous activity measurement and drug response in human cerebral organoids. (A) The adhesion surface between human cerebral organoids derived from Rett syndrome patients and CMOS-MEA is drawn with a black line. A heat map shows the number of firings detected in the spontaneous activity measurement for 153.6 seconds. (a) Vehicle, (b) 4-AP 100 µM. (B) Raster plot of 14,612 electrodes on the contact surface and histogram of the number of firings per 100 ms, (a) Vehicle, (b) 4-AP 100 µM. (C) Top row: 3-second 3D raster plot of the network burst indicated by the red line in B. The 3D raster plots represent time on the x-axis and electrodes to which the organoids are in contact in the Y–Z plane. The color scale of the plot indicates time from burst initiation. Bottom: Histogram of the number of firing electrodes per 5 ms for 3 seconds. (a) Vehicle, (b) 4-AP 100 µM. (B) Changes before and after administration of 4-AP, (a) total number of firings, (b) number of active electrodes, (c) normalized average firing frequency per electrode. (D) Changes before and after 4-AP 100 µM administration, (a) total number of firings, (b) number of active electrodes, (c) normalized average firing frequency per active electrode.

## Discussion

Using the world’s highest-spec HD-CMOS-MEA, which can sense a wide area of 5.5 mm × 5.9 mm with 236,880 electrodes (Figure 1), we recorded the propagation between the hippocampus and cerebral cortex with a sufficient S/N ratio (Figure 2). In the case of mouse brain slices, most of the brain regions can be measured simultaneously, and able to analyze the propagation between brain regions in detail. As shown in the voltage color map in Figure 2E and supplementary movie 1, it is a feature of our HD-CMOS-MEA that the tissue structure of the brain slice can be understood from the LFP sink and source. We found a relationship between sink and source in the apical dendrites and dorsal dendrite regions of the hippocampal pyramidal neuron (Figure 2C-g). However, sink and source appeared depending on the firing pattern, even in the same dendrite region. It has been reported that the sink and source patterns vary depending on the input^61^. By using our HD-CMOS-MEA, it will be possible to investigate the diversity of sinks and sources extensively.

Synchronous activity in the hippocampus and cortex was observed (Figure 2E). A report by Cappaert showed that the entorhinal cortex (EC) to hippocampus CA1 pathway and the EC to CA3 pathway were observed from the electrical activity of brain slices^62^, and it is thought that the hippocampal-cortico synchronous activity obtained in this study is the same result. Since the propagation of EC and hippocampal is observed in epilepsy, detailed time-series pattern measurement will advance research on the timing of EC-hippocampal synchronous activity. Additionally, through disease models and pharmacological tests, it is expected that epilepsy will be understood and applied to drug discovery and development.

Cognitive function in the brain involves synchronizing neural activity in the gamma frequency band. Cognitive deficits in schizophrenia are associated with abnormalities in the gamma-oscillation of cortical circuits^63^. Oscillations of high-frequency activity, known as SPW-R, are thought to facilitate communication between the hippocampus and the neocortex and promote long-term memory storage in cortical networks through strengthening intracortical connections^64^. It is also known that SPW-R transmission from the hippocampal dorsal CA1 to the granular retrosplenial cortex (gRSC) conducts via the subiculum^65^. SPW-R is also a marker of epilepsy, such as Dravet syndrome^66^, and in a model of schizophrenia^67^, SPW-R frequency, power, and frequency changes have been reported. In our HD-CMOS-MEA, we succeeded in capturing the propagation pattern of γ-oscillation, SPW-R propagation in the hippocampus, and propagation to the cortex (Figure 3). Propagation analysis of frequency characteristics using our HD-CMOS-MEA has the characteristic of being able to analyze over a wide area, and it is thought that it will bring new understandings on cognitive functions and neurological diseases, and for evaluation of convulsive toxicity of pharmaceuticals.

We also found that the electrical activity of the hiPS cell-derived cortical network can be analyzed on a cell-by-cell basis (Figure 4–6). Traditional MEA recordings also provide a means of generating electrophysiological activity from a large number of neurons. However, resolving the activity of individual neurons is challenging due to the poor spatial resolution of MEA data acquisition since the density of detectors is sparse on a cellular scale, which effectively renders each electrode a point detector. Despite this, it is possible to deconvolute the mixed signals of multiple neurons detected on each recording electrode of a MEA through the spike sorting analysis in which shape parameters of waveforms arising from a population of neurons are binned by similarity allowing for attribution of individual waveforms to a cell of origin^68, 69, 70, 71^. Parameters such as spike time tiling coefficient are used in inferring whether the two cells are functionally connected. Those bona fide connections thereby are considered to represent network activity among the neuron pairs^71, 72, 73, 74^. However, the results of spike sorting performed in this fashion come with the caveat that there is no “ground-truth” since it is impossible to know exactly how many individual neurons are being recorded. Signals arising from separate cells may be conflated as coming from the same point of origin or vice versa due to fluctuations in field potentials and propagation of potentials across the array^68, 69^. On the other hand, the CMOS-MEA provides high spatiotemporal-resolution electrical recording within a large sensor area at high electrode density, which allows for the acquisition of electrophysiological recordings at a subcellular resolution^75, 76, 77, 78, 79^. CMOS-MEA detects signals with high sensitivity and allows to pack electrodes to be very close to each other. Therefore, the spontaneous activity from an individual neuron could be recorded by multiple electrodes with high signal-to-noise ratios. The advanced spike sorting method^80^ achieves single-cell analysis for primary-neuron cultures on CMOS-MEA, and the network connectivity is detected between specified neuron pairs^15, 80^. However, this single-neuron analysis using CMOS-MEA is limited for signals only recorded from a preselected area up to about 1,000 electrodes, due to the circuit-design constraints, i.e., the little available area to realize high-performance circuits. Also, the network connection study is suboptimal compared to a whole-sample recording. Compared to the previous reports, we showed here a single-neuron analysis for hiPSC-derived neuron cultures on our HD-CMOS-MEA after a whole-sample recording with more than 59,220 electrodes, which largely improved the state-of-the-art.

The feature of increasing the NB to 4-AP and the feature of increasing the duration in a NB of PTX are the results reported in the conventional MEA measurement^19^. Owing to the parameters with significant differences were different between network analysis and single neuron analysis, it is considered that a more accurate analysis will be possible by analyzing them together. The Z score with other neurons calculated from the time-series data for each neuron is also considered an effective analysis method which reflects changes in synaptic connection strength.

4-AP acts as a K + channel blocker and Ca channel agonist, increasing the number of firings not only in excitatory neurons and inhibitory neurons, so it is thought that a significant increase in the rate of excitability binding was not observed (Figure 5 and 6). PTX, on the other hand, increased excitatory binding strength by blocking GABA-A receptors. It has been reported that PTX increases the rate of excitability binding and changes to regular activity^81^. Since the increase in regularity is reflected in the Z score, it is thought that the Z score increased because of the increase in excitatory synaptic connections (Figures 5 and 6). The decrease in ERC in Figure 6 reflects inhibitory connections. However, it is challenging to detect inhibitory binding as a numerical value by evaluating inhibitory binding only by Z score (Figure 5) since synchronous burst firing occurs. If only inhibitory binding is to be quantified, focusing on firings other than synchronous bursts may be effective.

Combining hiPSC-derived neurons with large-scale electrophysiological techniques, such as CMOS-MEAs, is thought to provide a powerful and scalable platform for neuropathy study^38, 82, 83^. However, such research is rarely reported, partly due to the lack of suitable metrics in downstream data analysis^84^. It is reported that, retigabine, an anticonvulsant, reduces neural activity in a dose-dependent manner from hiPSC-derived neuron cultures on CMOS-MEA, based on traditional network burst analysis^38^. However, the number and density of selected electrodes could be largely influenced by the result. A comparably large number of electrodes (1020 electrodes simultaneously in our report) could increase the reliability of extracted metrics. In this meaning, approaching the present study, a large-scale single-neuron analysis based on the big data acquisition by our HD-CMOS-MEA and the analysis of synaptic connection using Z score showed considerable potential to extract highly reproducible metrics for accurate assessment of drug effects on hiPSC-derived neuron cultures.

DRG neurons are composed of several types of neurons and glial cells with large heterogeneity. Traditionally, DRGs are categorized into three subtypes based on their morphology such as soma size and myelinated/unmyelinated fibers^85^. With the development of single-cell leveled analysis, the classification of DRG neurons was further subdivided into 17 clusters based on different gene expression patterns^86, 87, 88, 89, 90, 91^. However, one limitation of such large-scale single neuron classification based on similar transcriptomic profiles is the lack of direct functional correlates^92^. In the present study, we provide a strategy in identifying DRG single-cell classification based on neural electric activity, which has not been reported before. By using HD-CMOS-MEA, over 300 individual DRG neurons could be monitored simultaneously, which enable large-scale analysis and up to 10 subtypes of DRG neurons were classified by the frequency of spontaneous activity. Interestingly, we observed DRG neurons with a relatively high firing rate (close to 100 Hz), but at a low population (1% of whole identified DRG neurons). Neurons with high firing frequency were well identified in the cortical areas^93, 94^, but never in the peripheral nerves. A critical finding of this study is that we observed heterogeneous responses to TRP agonists in different subtypes of DRG neurons, which means that it is possible to identify particular functional responses in the specific DRG subtype. Furthermore, we would also like to classify DRG neuron subtypes based on morphology and single-cell transcriptomic expression in the future. Therefore, it is possible, in principle, to characterize the functional role of a specific gene isoform/ion channel/receptor in the relevant neuronal circuit to peripheral pain or toxicity.

Paclitaxel is widely used in treating some tumors, however it is known to cause peripheral neuropathy^95, 96^. A vitro study confirmed paclitaxel toxicity in cultured DRG neurons after 1 µM exposure for 24 h^97^. Using rat models, it is confirmed that TRPV1 upregulation contributes to paclitaxel-induced peripheral neuropathy^98, 99^. And Li’s group has shown that 12.5 µM paclitaxel treatment over 1 d could be significantly sensitive TRPV1 response to capsaicin burning in cultured DRG neurons^100^. In the present study, we have shown the same increase of TRPV1 sensitivity to capsaicin after paclitaxel treatment (at a lower concentration), possibly due to the high spatiotemporal resolution as proposed by our HD-CMOS-MEA, which allows the detecting of DRG neuronal activity at a single-cell level. Paclitaxel could also potentiate mechanosensor-mediated signal transduction, such as the piezo2 channel, in cultured cells^101^. Piezo2 has been confirmed to locate in rat peripheral sensory neurons, and increasing expression level of piezo2 is related to mechanical sensitivity in multiple pain conditions such as touch sensation or innoxious mechanic pain in rat models ^102, 103^. Therefore, it is considered that paclitaxel administration could advance mechanosensitive channels such as piezo2 in cultured DRG neurons, which induced higher neuronal sensitivity to vehicle administration, as observed in the present study.

Axonal parameters, such as axonal extension length and signal propagation velocity, can be used in investigating axonal dysfunction related to neurodegenerative diseases and drug testing^104^. In previous studies, microfluidic devices were used to allow axon elongation and signal propagation along axon was recorded by patch clamp or MEA^105, 106, 107, 108^. However, due to the very low amplitude of the axonal signals, these studies relied on repetitive electrical stimulation of an axonal segment to track signal propagation. CMOS-MEAs provide high temporal resolution, and therefore help trigger spontaneous signal propagation along the axonal arbors, such as in cultured primary hippocampus neurons and hiPSC-derived cortical neurons^15, 38, 109, 110^. Shimba’s group has also recorded signal conduction along myelinated DRG fibers under optical stimulation using a CMOS-MEA^111^. We recorded spontaneous axonal conduction dynamics in cultured DRG neurons in the present study using HD-CMOS-MEA. With the high spatiotemporal resolution provided by HD-CMOS-MEA, it has not necessary to compartmentalize DRG cultures using microfluidic, but detailed information including axonal pathway map and latency time, was directly shown. Since most peripheral nerves (particularly nociceptive C-fibers) are unmyelinated, this result is also considered to recapitulate in vivo mechanisms. Exactly, potential conduction velocities were found to be 1–2 m/s when measuring unmyelinated C-fibers in rats^112^, which shows the same order as our result.

Interestingly, a backward signal propagation initiated from the distal part of the axon was observed in DRG neurons. Such bidirectional signal conduction was also recently reported in the organotypic DRG explants^113^. A backward propagation action potential is wildly explored in the central nervous system, and is thought to provide necessary conditions for synaptic plasticity^114^. However, to our knowledge, no in vivo evidence exists of signal backpropagation in peripheral nerves, and the functional implications of such signal backpropagation in DRG neurons are still unclear.

Vincristine is a wildly used chemotherapeutic drug to treat a variety of cancers, but could induce serious peripheral neuropathy, which is the main factor restricting its clinical application^115^. The present study observed an initial increase in propagation velocity after 2 h vincristine treatment. It is reported that anticancer drugs could increase poration of membranes which leads to hyperexcitability at low, pre-toxic concentrations^116, 117^, that can explain the increased activity while the propagation velocity was reduced after 24 h and returned to the same level as before the vincristine treatment, while the propagation distance was further reduced. Indeed, several groups have reported that vincristine treatment did not influence nerve conduction velocity but could induce axonal degeneration in clinical studies^118, 119, 120^. This phenomenon corresponds to our findings. However, vincristine treatment was performed at a significantly lower concentration here than previous studies. It should also be noticed that several groups have shown conduction velocity reduction under high-dose vincristine treatment^121, 122^, probably due to the asymmetric soma size/diameter in different DRG subtypes, as mentioned above. We would like to explore the relationship between the conduction velocity and the concentration or treatment time of vincristine in the future, to further understand the mechanisms related to chemotherapy-induced peripheral neuropathy. Axonal conduction velocity measurement with HD-CMOS-MEA has the characteristic of being able to capture functional changes at low doses.

Human brain organoids serve as samples for the elucidation of neurological diseases, drug screening, and drug toxicity prediction research from the perspective of *in vitro* to *in vivo* extrapolation (IVIVE). The functional evaluation of brain organoids is mainly performed by MEA measurement, and it is reported that the electrophysiological characteristics of brain organoids which were obtained by MEA measurement are correlated with the maturation of neural networks over time revealed by scRNA-seq.^52^, demonstrating the advantages of having a 3D structure and the effectiveness of MEA measurement. In addition, drug screening using the electrophysiological characteristics of brain organoids obtained by MEA measurement has been proposed^123^. It has been reported that the promising effects of NitroSynapsin for the treatment of AD were captured using human brain organoids^124^. These reports indicate that human brain organoids are excellent samples for drug screening and neurological disease research in extrapolation to Vivo. By analyzing low-frequency signals obtained by MEA measurement of cerebral organoids, we evaluated the anticonvulsant properties of drugs and the effects of antiepileptic drugs. We reported that the results were effective for *in vivo* extrapolation^125^. Regarding brain organoid measurement using CMOS-MEA, it has been reported that CMOS-MEA measurements with 1,024 recording electrodes can evaluate the effects of drugs on large-scale and local neural networks^126^. However, the activity data of brain organoids obtained by CMOS-MEA measurement is a huge amount of big data, and the analysis method is highly complicated. It is expected that the establishment of an analysis method will be a breakthrough for further advancements. In this study, we measured electrical activity at the single-cell level with 7,260 electrodes over a wide range. In addition, detecting a phenomenon in which activity frequency increases in 4-AP administration showed that it is an assessment method that can be used to predict the seizure liability and drug efficacy of pharmaceuticals. The increased activity frequency for 4-AP was also observed in hiPSC-derived neurons and brain slices^19, 127, 128^, and showed a similar tendency. An expansion of the synchronous activity range (Figure 11C) was observed, and it was found to have a feature that allows analysis of drug response based on more detailed propagation patterns. Although the correlation of electrical activity with the structure of brain organoids required to be analyzed in future, the HD-CMOS-MEA measurement of brain organoids may be helpful for drug screening and functional evaluation of human brain diseases using diseased brain organoids.

We present a detailed and single-cell-level neural activity analysis platform for brain slices, human iPS cell-derived cortical networks, peripheral neurons, and human brain organoids. The detailed analysis of neural activity at the single-cell level using our CMOS-MEA provides a new platform for understanding the basic mechanisms of brain circuits *in vitro* and *ex vivo*, exploring human neurological diseases, drug discovery, and compound toxicity assessment.

## Methods

### Brain slices

C57BL/6NCrSlc mice (Six weeks old) were obtained from Japan SLC. Inc. Slices of mice brain were prepared as 300 μm using a slicer (NeoLinearSlicer MT, Dosaka EM). The study was approved by Tohoku Institute of Technology Animal Care and User Committee. Brain slices were placed on CMOS-MEA coated with 0.1% polyethylenimin (Sigma). During the experiments, the CMOS-MEA were perfused with artificial cerebrospinal fluid (in [mM]: NaCl (KCH5696, FUJIFILM) 124, KCl (160-03555, Wako) 3, NaH2PO4 (169-04245, Wako) 1.25, MgSO4(131-00405, Wako) 0.1, Glucose(044-00695, Wako) 10, NaHCO3(191-01305, Wako) 26, CaCl2(039-00475, Wako) 14.5 with carbogen (95% O2, 5% CO2) at a flow rate of 1.5 mL/min using a perfusion device [(Gilson, Inc.) Mini Pulse Pump Ⅲ MP-4]. The local field potential (LFP) of each brain region was measured at 1 kHz sampling rate with 236,880 electrodes simultaneously.

### Culture of human iPSC-derived cortical neurons

Before cell seeding, the surface of the Sony CMOS-MEA plate was coated sequentially by Cellmatrix collagen type I-C solution (637-00773, Nitta Gelatin), Poly-D-lysine solution (P7405, Sigma-Aldrich), and iMatrix 511 laminin solution (892019, Matrixome), each for 1 h. Cryopreserved hiPSC-derived cortical neurons (XCL-1 Neurons, XCell Science) were thawed and suspended in Neuron Medium (XCS-NM-001-M100-1P, XCell Science). For dispersed culture, approximately 7.0 × 10^4^ cells (8.0 × 10^5^ cells/cm^2^) in 15 µL neuron medium were seeded directly in the middle of the Sony CMOS-MEA plate at the location of the electrode array. After 30 min, 1 mL of neural maturation basal medium (NM-001-BM100, XCell Science Inc., USA) supplemented with neuron maturation supplement A (NM-001-SA100, XCell Science Inc., USA) and 100 U/mL penicillin/streptomycin (168–23191, Wako) was applied. After one week, the medium was replaced with 1 mL of BrainPhys neuronal medium containing SM1 neuronal supplement (ST-05792, STEMCELL technologies) and Human iPSC-derived mature astrocytes (XCL-1 mature astrocytes, AR-001-1V, XCell Science) were seeded at approximately 7.0 × 10^4^ cells in the Sony CMOS-MEA plate. Following this, half the volume of the medium was replaced twice per week.

### Culture of DRG neurons

DRG neurons were harvested and cultured as described previously^129^. Briefly, DRG neurons were collected from a 10-week-old male Wistar Rats. Firstly, the rats were asphyxiated with isoflurane and then decapitated. DRGs were harvested from the vertebral column, and the sensory neurons were dissociated by mechanical agitation after incubation for 2 h with collagenase type III (CLS3, Worthington) at 37°C. Then, the cells were washed with Hank’s balanced salt solution and further dissociated with trypsin type I (T8003, Sigma-Aldrich). After cell counting, approximately 5.0 × 10^4^ cells (6.0 × 10^5^ cells/cm^2^) in 15 µL BrainPhys Neuronal Medium were seeded directly in the middle of the Sony CMOS-MEA plate. After 30 min, 1 mL of BrainPhys neuronal medium was applied. The next day, the medium was replaced with 1 mL of serum-free medium containing 10 µm uridine and 10 µm 2′-deoxy-5-fluorouridine kept for 3 days to suppress the proliferation of glial cells. Afterwards, the medium was changed back to 1 mL BrainPhys neuronal medium, and half the volume of the medium was replaced twice per week.

### Culture of human cerebral organoids

Human-derived iPSCs (HPS3036, Disease-specific iPS cell line derived from a patient: Rett syndrome) were obtained from the Institute of Physical and Chemical Research. Briefly, iPSCs were cultured by StemFit (AK02 N, Ajinomoto) and were collected using Gentle Cell Dissociation Reagent (ST-07174, STEMCELL Technologies) when cells were confluent on six-well dishes. The collected cells were centrifuged for 5 mins at 800 rpm at room temperature. After discarding the supernatant, 1 ml EB seeding medium (EB formation medium added 10 mM Y-27632) was added and the cell pellet was resuspended. iPSCs were cultured at 9.0 × 103 cells/wel in 96 wells using the EB seeding medium. After 2 and 4 days, 100 μl EB seeding medium was added per well. On day five, organoids were observed and spherical form samples were selected. EB seeding medium was substituted with an induction medium and incubated for 2 days. Organoids were embedded in Matrigel (354,277, Corning) and incubated in an expansion medium for 3 days. The expansion medium was substituted with a maturation medium, and the organoids were incubated in an orbital shaker (COSH6, AS ONE Corporation). Organoids were maintained in the maturation medium for 3 months, and the medium replenishment was performed every 3–4 days. After 3 months, the culture medium was changed to Brain Phys (ST-05792, STEMCELL Technologies). The Medium of the STEMdiff Cerebral Organoid Kit (ST-08570, STEMCELL Technologies) was used for organoid formation.

### Immunocytochemistry

Sample cultures were fixed with 4% paraformaldehyde in PBS on ice (4°C) for 10 mins. Fixed cells were incubated with 0.2% Triton-X-100 in PBS for 5 min, then with preblock buffer (0.05% Triton-X-100 and 5% goat serum in PBS) at 4°C for 1 h, and finally with preblock buffer containing a specific primary antibody (1:1,000) at 4°C for 24 h. The primary antibodies used were rabbit anti-MAP2 (ab281588, abcom) and mouse anti-β-tubulin III (T8578, Sigma-Aldrich). Then, the samples were incubated with the appropriate secondary antibody (anti-rabbit 488 Alexa Fluor, ab150077, Abcam or anti-mouse 488 Alexa Fluor, ab150113, Abcam, 1:1,000 in preblock buffer) for 1 h at room temperature. For hiPSC-derived cortical neuron sample, cell nuclei were counterstained with 1 μg/mL Hoechst 33258 (H341, DOJINDO) for 1 h at room temperature. Stained cultures were washed twice with preblock buffer (5 min/wash) and rinsed twice with PBS. The immunolabeling was visualized by using a confocal microscope (Eclipse Ni, Nikon). Image intensity was adjusted using the ImageJ software (NIH).

### Pharmacological tests

For hiPSC-derived cortical neuron samples, spontaneous activities were recorded before treatment and after the cumulative addition to the culture medium of one of the following compounds: 4-AP (10 and 30 µM; 016-02781, Wako), picrotoxin (1 and 10 µM; 2800471, Nacalai tesque), and AP-5 (25 µM; ab120003, Abcam) followed by CNQX (30 µM; 032-23121, Wako). All chemicals were dissolved in DMSO (0.1%), used as a vehicle.

For DRG samples, spontaneous activities were recorded before and after the cumulative addition of capsaicin (030-11353, Wako) at 1, 10, and 100 nM to the culture medium. Then the medium was replaced with a culture medium containing 10 nM paclitaxel (161-28164, Wako). After 24 h culture, spontaneous activities were rerecorded before and after capsaicin treatment.

For conduction velocity analysis experiments using DRG samples, spontaneous activities were recorded before, 2 h after, and 24 h after adding 3 nM vincristine (220-02301, Wako) to the culture medium.

In the recordings and drug administration, the cultures were kept at 37°C under a 5% CO2 atmosphere.

### Extracellular recording

Spontaneous extracellular field potentials were acquired at 37 °C under a 5% CO2 atmosphere using a HD-CMOS-MEA system (Sony semiconductor solutions) at a sampling rate of 1–70 kHz/channel^57^. The spikes in the acquired data were detected using a 100-Hz high-pass filter. The measurement time was controlled by a software (Sony semiconductor solutions), and all the raw data was saved on the PC.

### Frequency analysis

A bandpass filter using the MATLAB function “filtfilt,” a zero-phase filtering, was used to extract each frequency component of the raw data obtained by brain slice experiments.

### Spike detection

Spikes at each electrode of hiPSC-derived cortical neuron samples were detected using a voltage threshold of ±80 μV. In detecting single-cell firing, electrodes with identical firing patterns were combined. Specifically, the firing frequency histogram of 1 ms bins for each electrode was calculated, and electrodes with a cosine similarity of 0.3 or more in histograms between adjacent electrodes were combined as electrodes of the same neuron. Spikes at each electrode of DRG samples in capsaicin response were detected using a voltage threshold of ±100 µV. The spikes at each electrode of organoid samples were detected using a voltage threshold of ±200 μV. Spike detection and soma identification were performed using MATLAB (Mathworks Natick, MA).

### Burst analysis

Network bursts in hiPSC-derived cortical neuron samples and organoid samples were detected using a 4-step method^130^ similar to the synchronous burst detection method in conventional MEAs. Briefly, inter-spike interval (ISI) is calculated from time-stamped data from multiple neurons, and bursts are detected by setting thresholds for the ISI threshold, inter-burst interval threshold, and the number of spikes within a burst.

Single neuron burst at hiPSC-derived cortical neuron samples were detected using Poisson-Surprise (PS) method^131, 132^. Divide the timestamp data into groups of firings with intervals shorter than the average firing interval, and extract the firing group with the maximum PS value for each firing group. Here we assume that each neuron fires according to the Poisson distribution. A threshold was set for the PS value of the extracted firing group and the number of firings within the burst, and single-cell bursts were detected. Network burst analysis and single neuron burst analysis used MATLAB in calculating the parameters.

### Connection analysis

We applied pairwise synchronous analysis using MATLAB to investigate the temporal correlation of spikes between individual neurons in hiPSC-derived cortical neuron samples.

### Capsaicin-induced response analysis

Pairwise Euclidean distances between each cell, which were defined by a histogram vector of the number of firings for 10 seconds after capsaicin drug administration, were calculated. Dendrogram analysis was used in classifying neurons into five firing types based on firing pattern similarity in drug response. MATLAB was used for the series of analyses.

### Statistical analysis

Multiple group comparisons were performed using one-way ANOVA followed by Dunnett’s test or Holm’s test were used to calculate the significant difference between each concentration.

### Data availability

The data and scripts that support the findings of this study are available from the corresponding author upon reasonable request.

## Acknowledgments

This study was supported by the grant of collaborative project with Sony semiconductor solutions Inc and Adaptable and Seamless Technology transfer Program through Target-driven R&D (A-STEP) from Japan Science and Technology Agency (JST) Grant Number JPMJTR20UP.

## Author contributions

I.S. designed experiments; X.H., S.N., M.S., and N.N. conducted experiments; N.M., S.N., X.H., and Y.I. analyzed the data; I.S., N.M., and X.H. wrote the manuscript; I.S. supervised the project.

## References

1. Hong G, Lieber CM. Novel electrode technologies for neural recordings. Nature Reviews Neuroscience 20, 330–345 (2019).

2. Chen ZS, Pesaran B. Improving scalability in systems neuroscience. Neuron 109, 1776–1790 (2021).

3. Paulk AC, et al. Large-scale neural recordings with single neuron resolution using Neuropixels probes in human cortex. Nature Neuroscience 25, 252–263 (2022).

4. Meisenhelter S, Rutishauser U. Probing the human brain at single-neuron resolution with high-density cortical recordings. Neuron 110, 2353–2355 (2022).

5. Jun JJ, et al. Fully integrated silicon probes for high-density recording of neural activity. Nature 551, 232–236 (2017).

6. Raducanu BC, et al. Time Multiplexed Active Neural Probe with 1356 Parallel Recording Sites (2017).

7. Rios G, Lubenov EV, Chi D, Roukes ML, Siapas AG. Nanofabricated Neural Probes for Dense 3-D Recordings of Brain Activity. Nano Letters 16, 6857–6862 (2016).

8. Chung JE, et al. High-Density, Long-Lasting, and Multi-region Electrophysiological Recordings Using Polymer Electrode Arrays. Neuron 101, 21–31.e25 (2019).

9. Angotzi GN, et al. SiNAPS: An implantable active pixel sensor CMOS-probe for simultaneous large-scale neural recordings. Biosensors and Bioelectronics 126, 355–364 (2019).

10. Steinmetz NA, et al. Neuropixels 2.0: A miniaturized high-density probe for stable, long-term brain recordings. Science 372, eabf4588 (2021).

11. Fu T-M, Hong G, Zhou T, Schuhmann TG, Viveros RD, Lieber CM. Stable long-term chronic brain mapping at the single-neuron level. Nature Methods 13, 875–882 (2016).

12. Fiáth R, et al. Fine-scale mapping of cortical laminar activity during sleep slow oscillations using high-density linear silicon probes. Journal of Neuroscience Methods 316, 58–70 (2019).

13. Yuste R. From the neuron doctrine to neural networks. Nature Reviews Neuroscience 16, 487–497 (2015).

14. Yuan X, Emmenegger V, Obien MEJ, Hierlemann A, Frey U. Dual-mode Microelectrode Array Featuring 20k Electrodes and High SNR for Extracellular Recording of Neural Networks. IEEE Biomedical Circuits and Systems Conference : healthcare technology : [proceedings] IEEE Biomedical Circuits and Systems Conference 2018, (2019).

15. Yuan X, et al. Versatile live-cell activity analysis platform for characterization of neuronal dynamics at single-cell and network level. Nature Communications 11, 4854 (2020).

16. Takahashi K, Yamanaka S. Induction of Pluripotent Stem Cells from Mouse Embryonic and Adult Fibroblast Cultures by Defined Factors. Cell 126, 663–676 (2006).

17. Grainger AI, King MC, Nagel DA, Parri HR, Coleman MD, Hill EJ. In vitro Models for Seizure-Liability Testing Using Induced Pluripotent Stem Cells. Frontiers in neuroscience 12, (2018).

18. Black BJ, Atmaramani R, Pancrazio JJ. Spontaneous and Evoked Activity from Murine Ventral Horn Cultures on Microelectrode Arrays. Frontiers in Cellular Neuroscience 11, (2017).

19. Bradley JA, Luithardt HH, Metea MR, Strock CJ. In Vitro Screening for Seizure Liability Using Microelectrode Array Technology. Toxicological Sciences 163, 240–253 (2018).

20. Kreir M, et al. Do in vitro assays in rat primary neurons predict drug-induced seizure liability in humans? Toxicology and Applied Pharmacology 346, 45–57 (2018).

21. Toivanen M, et al. Optimised PDMS Tunnel Devices on MEAs Increase the Probability of Detecting Electrical Activity from Human Stem Cell-Derived Neuronal Networks. Frontiers in neuroscience 11, (2017).

22. Matsuda N, Odawara A, Kinoshita K, Okamura A, Shirakawa T, Suzuki I. Raster plots machine learning to predict the seizure liability of drugs and to identify drugs. Scientific reports 12, 2281 (2022).

23. Obien MEJ, Deligkaris K, Bullmann T, Bakkum DJ, Frey U. Revealing neuronal function through microelectrode array recordings. Frontiers in neuroscience 8, (2015).

24. Odawara A, Saitoh Y, Alhebshi AH, Gotoh M, Suzuki I. Long-term electrophysiological activity and pharmacological response of a human induced pluripotent stem cell-derived neuron and astrocyte co-culture. Biochemical and Biophysical Research Communications 443, 1176–1181 (2014).

25. Odawara A, Katoh H, Matsuda N, Suzuki I. Physiological maturation and drug responses of human induced pluripotent stem cell-derived cortical neuronal networks in long-term culture. Scientific reports 6, 26181 (2016).

26. Ojima A, Miyamoto N. [Method for MEA Data Analysis of Drug-treated Rat Primary Neurons and Human iPSC-derived Neurons to Evaluate the Risk of Drug-induced Seizures]. Yakugaku Zasshi 138, 823–828 (2018).

27. Kayama T, Suzuki I, Odawara A, Sasaki T, Ikegaya Y. Temporally coordinated spiking activity of human induced pluripotent stem cell-derived neurons co-cultured with astrocytes. Biochemical and Biophysical Research Communications 495, 1028–1033 (2018).

28. Ryynänen T, Toivanen M, Salminen T, Ylä-Outinen L, Narkilahti S, Lekkala J. Ion Beam Assisted E-Beam Deposited TiN Microelectrodes—Applied to Neuronal Cell Culture Medium Evaluation. Frontiers in neuroscience 12, (2018).

29. Sasaki T, Suzuki I, Yokoi R, Sato K, Ikegaya Y. Synchronous spike patterns in differently mixed cultures of human iPSC-derived glutamatergic and GABAergic neurons. Biochemical and Biophysical Research Communications 513, 300–305 (2019).

30. Shirakawa T, Suzuki I. Approach to Neurotoxicity using Human iPSC Neurons: Consortium for Safety Assessment using Human iPS Cells. Current pharmaceutical biotechnology 21, 780–786 (2020).

31. Ishibashi Y, Odawara A, Kinoshita K, Okamura A, Shirakawa T, Suzuki I. Principal Component Analysis to Distinguish Seizure Liability of Drugs in Human iPS Cell-Derived Neurons. Toxicological Sciences 184, 265–275 (2021).

32. Ichise E, et al. Impaired neuronal activity and differential gene expression in STXBP1 encephalopathy patient iPSC-derived GABAergic neurons. Human Molecular Genetics 30, 1337–1348 (2021).

33. Ghatak S, et al. NitroSynapsin ameliorates hypersynchronous neural network activity in Alzheimer hiPSC models. Molecular psychiatry 26, 5751–5765 (2021).

34. Klein Gunnewiek TM, et al. Mitochondrial dysfunction impairs human neuronal development and reduces neuronal network activity and synchronicity.). bioRxiv (2019).

35. Linda K, et al. Imbalanced autophagy causes synaptic deficits in a human model for neurodevelopmental disorders. Autophagy 18, 423–442 (2022).

36. Frega M, et al. Neuronal network dysfunction in a model for Kleefstra syndrome mediated by enhanced NMDAR signaling. Nat Commun 10, 4928 (2019).

37. Que Z, et al. Hyperexcitability and Pharmacological Responsiveness of Cortical Neurons Derived from Human iPSCs Carrying Epilepsy-Associated Sodium Channel Nav1.2-L1342P Genetic Variant. The Journal of Neuroscience 41, 10194 (2021).

38. Ronchi S, et al. Electrophysiological Phenotype Characterization of Human iPSC-Derived Neuronal Cell Lines by Means of High-Density Microelectrode Arrays. Advanced Biology 5, 2000223 (2021).

39. Simkin D, et al. Dyshomeostatic modulation of Ca(2+)-activated K(+) channels in a human neuronal model of KCNQ2 encephalopathy. eLife 10, (2021).

40. Wainger Brian J, et al. Intrinsic Membrane Hyperexcitability of Amyotrophic Lateral Sclerosis Patient-Derived Motor Neurons. Cell Reports 7, 1–11 (2014).

41. Odawara A, Shibata M, Ishibashi Y, Nagafuku N, Matsuda N, Suzuki I. In Vitro Pain Assay Using Human iPSC-Derived Sensory Neurons and Microelectrode Array. Toxicological sciences : an official journal of the Society of Toxicology 188, 131–141 (2022).

42. Xiong C, et al. Human Induced Pluripotent Stem Cell Derived Sensory Neurons are Sensitive to the Neurotoxic Effects of Paclitaxel. Clinical and translational science 14, 568–581 (2021).

43. Schinke C, et al. Modeling chemotherapy induced neurotoxicity with human induced pluripotent stem cell (iPSC) -derived sensory neurons. Neurobiology of disease 155, 105391 (2021).

44. Cunningham GM, et al. The impact of SBF2 on taxane-induced peripheral neuropathy. PLoS genetics 18, e1009968 (2022).

45. Vojnits K, Mahammad S, Collins TJ, Bhatia M. Chemotherapy-Induced Neuropathy and Drug Discovery Platform Using Human Sensory Neurons Converted Directly from Adult Peripheral Blood. Stem cells translational medicine 8, 1180–1191 (2019).

46. Wang M, et al. Mechanisms of peripheral neurotoxicity associated with four chemotherapy drugs using human induced pluripotent stem cell-derived peripheral neurons. Toxicology in vitro : an international journal published in association with BIBRA 77, 105233 (2021).

47. Kathuria A, et al. Transcriptomic Landscape and Functional Characterization of Induced Pluripotent Stem Cell–Derived Cerebral Organoids in Schizophrenia. JAMA Psychiatry 77, 745–754 (2020).

48. Trujillo CA, et al. Pharmacological reversal of synaptic and network pathology in human MECP2-KO neurons and cortical organoids. EMBO Molecular Medicine 13, e12523 (2021).

49. Sun G, et al. Modeling Human Cytomegalovirus-Induced Microcephaly in Human iPSC-Derived Brain Organoids. Cell Reports Medicine 1, 100002 (2020).

50. Shin H, Jeong S, Lee J-H, Sun W, Choi N, Cho I-J. 3D high-density microelectrode array with optical stimulation and drug delivery for investigating neural circuit dynamics. Nature Communications 12, 492 (2021).

51. Yao H, et al. Methadone interrupts neural growth and function in human cortical organoids. Stem Cell Research 49, 102065 (2020).

52. Fair SR, et al. Electrophysiological Maturation of Cerebral Organoids Correlates with Dynamic Morphological and Cellular Development. Stem Cell Reports 15, 855–868 (2020).

53. Bautista DM, et al. The menthol receptor TRPM8 is the principal detector of environmental cold. Nature 448, 204–208 (2007).

54. Jeske NA, et al. A-kinase anchoring protein mediates TRPV1 thermal hyperalgesia through PKA phosphorylation of TRPV1. Pain 138, 604–616 (2008).

55. McNamara CR, et al. TRPA1 mediates formalin-induced pain. Proceedings of the National Academy of Sciences 104, 13525–13530 (2007).

56. Samanta A, Hughes TET, Moiseenkova-Bell VY. Transient Receptor Potential (TRP) Channels. In: Membrane Protein Complexes: Structure and Function (eds Harris JR, Boekema EJ). Springer Singapore (2018).

57. Kato Y, et al. High-Density and Large-Scale MEA System Featuring 236,880 Electrodes at 11.72μm Pitch for Neuronal Network Analysis. In: 2020 IEEE Symposium on VLSI Circuits) (2020).

58. de Curtis M, Avanzini G. Interictal spikes in focal epileptogenesis. Progress in Neurobiology 63, 541–567 (2001).

59. Avoli M, de Curtis M. GABAergic synchronization in the limbic system and its role in the generation of epileptiform activity. Progress in Neurobiology 95, 104–132 (2011).

60. In: *Jasper’s Basic Mechanisms of the Epilepsies* (eds Noebels JL, Avoli M, Rogawski MA, Olsen RW, Delgado-Escueta AV). National Center for Biotechnology Information (US) Copyright © 2012, Michael A Rogawski, Antonio V Delgado-Escueta, Jeffrey L Noebels, Massimo Avoli and Richard W Olsen. (2012).

61. Herreras O. Local Field Potentials: Myths and Misunderstandings. Frontiers in Neural Circuits 10, (2016).

62. Cappaert NLM, Lopes da Silva FH, Wadman WJ. Spatio-temporal dynamics of theta oscillations in hippocampal–entorhinal slices. Hippocampus 19, 1065–1077 (2009).

63. Cho RY, Konecky RO, Carter CS. Impairments in frontal cortical γ synchrony and cognitive control in schizophrenia. Proceedings of the National Academy of Sciences 103, 19878–19883 (2006).

64. Buzsáki G. Hippocampal sharp wave-ripple: A cognitive biomarker for episodic memory and planning. Hippocampus 25, 1073–1188 (2015).

65. Nitzan N, et al. Propagation of hippocampal ripples to the neocortex by way of a subiculum-retrosplenial pathway. Nature Communications 11, 1947 (2020).

66. Cheah CS, Lundstrom BN, Catterall WA, Oakley JC. Impairment of Sharp-Wave Ripples in a Murine Model of Dravet Syndrome. The Journal of Neuroscience 39, 9251 (2019).

67. Suh J, Foster David J, Davoudi H, Wilson Matthew A, Tonegawa S. Impaired Hippocampal Ripple-Associated Replay in a Mouse Model of Schizophrenia. Neuron 80, 484–493 (2013).

68. Buzsáki G, Anastassiou CA, Koch C. The origin of extracellular fields and currents— EEG, ECoG, LFP and spikes. Nature reviews neuroscience 13, 407–420 (2012).

69. Gibson SP. Neural spike sorting in hardware: From theory to practice. University of California, Los Angeles (2012).

70. Kyung Hwan K, Sung June K. Neural spike sorting under nearly 0-dB signal-to-noise ratio using nonlinear energy operator and artificial neural-network classifier. IEEE Transactions on Biomedical Engineering 47, 1406–1411 (2000).

71. Negri J, Menon V, Young-Pearse TL. Assessment of Spontaneous Neuronal Activity In Vitro Using Multi-Well Multi-Electrode Arrays: Implications for Assay Development. eNeuro 7, (2020).

72. Slomowitz E, et al. Interplay between population firing stability and single neuron dynamics in hippocampal networks. eLife 4, e04378 (2015).

73. Vincent K, Tauskela JS, Mealing GA, Thivierge J-P. Altered Network Communication Following a Neuroprotective Drug Treatment. PLOS ONE 8, e54478 (2013).

74. Cutts CS, Eglen SJ. Detecting Pairwise Correlations in Spike Trains: An Objective Comparison of Methods and Application to the Study of Retinal Waves. The Journal of Neuroscience 34, 14288 (2014).

75. Tsai D, Sawyer D, Bradd A, Yuste R, Shepard KL. A very large-scale microelectrode array for cellular-resolution electrophysiology. Nature Communications 8, 1802 (2017).

76. Eversmann B, et al. CMOS sensor array for electrical imaging of neuronal activity. In: 2005 IEEE International Symposium on Circuits and Systems (ISCAS)) (2005).

77. Berdondini L, et al. Active pixel sensor array for high spatio-temporal resolution electrophysiological recordings from single cell to large scale neuronal networks. Lab on a chip 9, 2644–2651 (2009).

78. Ballini M, et al. A 1024-Channel CMOS Microelectrode Array With 26,400 Electrodes for Recording and Stimulation of Electrogenic Cells In Vitro. IEEE Journal of Solid-State Circuits 49, 2705–2719 (2014).

79. Dragas J, et al. In Vitro Multi-Functional Microelectrode Array Featuring 59 760 Electrodes, 2048 Electrophysiology Channels, Stimulation, Impedance Measurement, and Neurotransmitter Detection Channels. IEEE Journal of Solid-State Circuits 52, 1576–1590 (2017).

80. Yger P, et al. A spike sorting toolbox for up to thousands of electrodes validated with ground truth recordings in vitro and in vivo. eLife 7, e34518 (2018).

81. Hablitz JJ. Picrotoxin-induced epileptiform activity in hippocampus: role of endogenous versus synaptic factors. Journal of neurophysiology 51, 1011–1027 (1984).

82. Sundberg M, et al. 16p11.2 deletion is associated with hyperactivation of human iPSC-derived dopaminergic neuron networks and is rescued by RHOA inhibition in vitro. Nature Communications 12, 2897 (2021).

83. Lopez CM, et al. A Multimodal CMOS MEA for High-Throughput Intracellular Action Potential Measurements and Impedance Spectroscopy in Drug-Screening Applications. IEEE Journal of Solid-State Circuits 53, 3076–3086 (2018).

84. McCready FP, Gordillo-Sampedro S, Pradeepan K, Martinez-Trujillo J, Ellis J. Multielectrode Arrays for Functional Phenotyping of Neurons from Induced Pluripotent Stem Cell Models of Neurodevelopmental Disorders (2022).

85. Kandel ER, Schwartz JH, Jessell TM, Mack S, Dodd J. *Principles of neural science*, 4th ed edn. McGraw-Hill, Health Professions Division (2000).

86. Li C-L, et al. Somatosensory neuron types identified by high-coverage single-cell RNA-sequencing and functional heterogeneity. Cell Research 26, 83–102 (2016).

87. Nguyen MQ, Le Pichon CE, Ryba N. Stereotyped transcriptomic transformation of somatosensory neurons in response to injury. eLife 8, e49679 (2019).

88. Renthal W, et al. Transcriptional Reprogramming of Distinct Peripheral Sensory Neuron Subtypes after Axonal Injury. Neuron 108, 128–144.e129 (2020).

89. Sharma N, Flaherty K, Lezgiyeva K, Wagner DE, Klein AM, Ginty DD. The emergence of transcriptional identity in somatosensory neurons. Nature 577, 392–398 (2020).

90. Usoskin D, et al. Unbiased classification of sensory neuron types by large-scale single-cell RNA sequencing. Nature Neuroscience 18, 145–153 (2015).

91. Zeisel A, et al. Molecular Architecture of the Mouse Nervous System. Cell 174, 999–1014.e1022 (2018).

92. Nelson SB, Sugino K, Hempel CM. The problem of neuronal cell types: a physiological genomics approach. Trends in Neurosciences 29, 339–345 (2006).

93. Connors BW, Gutnick MJ. Intrinsic firing patterns of diverse neocortical neurons. Trends in Neurosciences 13, 99–104 (1990).

94. McCormick DA, Connors BW, Lighthall JW, Prince DA. Comparative electrophysiology of pyramidal and sparsely spiny stellate neurons of the neocortex. Journal of neurophysiology 54, 782–806 (1985).

95. Hagiwara H, Sunada Y. Mechanism of taxane neurotoxicity. Breast Cancer 11, 82–85 (2004).

96. Chaudhry V, Rowinsky EK, Sartorius SE, Donehower RC, Cornblath DR. Peripheral neuropathy from taxol and cisplatin combination chemotherapy: Clinical and electrophysiological studies. Annals of Neurology 35, 304–311 (1994).

97. Scuteri A, et al. Paclitaxel Toxicity in Post-mitotic Dorsal Root Ganglion (DRG) Cells. Anticancer Research 26, 1065 (2006).

98. Hara T, et al. Effect of paclitaxel on transient receptor potential vanilloid 1 in rat dorsal root ganglion. PAIN® 154, 882–889 (2013).

99. Kamata Y, et al. Paclitaxel Induces Upregulation of Transient Receptor Potential Vanilloid 1 Expression in the Rat Spinal Cord (2020).

100. Li Y, et al. The Cancer Chemotherapeutic Paclitaxel Increases Human and Rodent Sensory Neuron Responses to TRPV1 by Activation of TLR4. The Journal of Neuroscience 35, 13487 (2015).

101. Chang W, Gu JG. Role of microtubules in Piezo2 mechanotransduction of mouse Merkel cells. Journal of neurophysiology 124, 1824–1831 (2020).

102. Shin SM, et al. Piezo2 mechanosensitive ion channel is located to sensory neurons and nonneuronal cells in rat peripheral sensory pathway: implications in pain. Pain 162, (2021).

103. Tu H, Zhang D, Li Y-L. Cellular and Molecular Mechanisms Underlying Arterial Baroreceptor Remodeling in Cardiovascular Diseases and Diabetes. Neuroscience Bulletin 35, 98–112 (2019).

104. Millecamps S, Julien J-P. Axonal transport deficits and neurodegenerative diseases. Nature Reviews Neuroscience 14, 161–176 (2013).

105. Kramer L, et al. Modeling chemotherapy-induced peripheral neuropathy using a nerve-on-a-chip microphysiological system. ALTEX – Alternatives to animal experimentation 37, 350–364 (2020).

106. Black BJ, et al. Emerging neurotechnology for antinoceptive mechanisms and therapeutics discovery. Biosensors and Bioelectronics 126, 679–689 (2019).

107. Tsantoulas C, Farmer C, Machado P, Baba K, McMahon SB, Raouf R. Probing Functional Properties of Nociceptive Axons Using a Microfluidic Culture System. PLOS ONE 8, e80722 (2013).

108. Rumsey JW, et al. Classical Complement Pathway Inhibition in a “Human-On-A-Chip” Model of Autoimmune Demyelinating Neuropathies. Advanced Therapeutics 5, 2200030 (2022).

109. Bakkum DJ, et al. Tracking axonal action potential propagation on a high-density microelectrode array across hundreds of sites. Nature Communications 4, 2181 (2013).

110. Buccino AP, Yuan X, Emmenegger V, Xue X, Gänswein T, Hierlemann A. An automated method for precise axon reconstruction from recordings of high-density micro-electrode arrays. Journal of neural engineering 19, 026026 (2022).

111. Shimba K, Asahina T, Sakai K, Kotani K, Jimbo Y. Recording Saltatory Conduction Along Sensory Axons Using a High-Density Microelectrode Array. Frontiers in neuroscience 16, (2022).

112. Zotova EG, Arezzo JC. NON-INVASIVE EVALUATION OF NERVE CONDUCTION IN SMALL DIAMETER FIBERS IN THE RAT. Physiology journal 2013, (2013).

113. Mateus JC, et al. Bidirectional flow of action potentials in axons drives activity dynamics in neuronal cultures. Journal of neural engineering 18, 066045 (2021).

114. Stuart G, Spruston N, Sakmann B, Häusser M. Action potential initiation and backpropagation in neurons of the mammalian CNS. Trends in Neurosciences 20, 125–131 (1997).

115. Li G-z, et al. Vincristine-induced peripheral neuropathy: A mini-review. NeuroToxicology 81, 161–171 (2020).

116. Zhang H, Dougherty PM. Enhanced Excitability of Primary Sensory Neurons and Altered Gene Expression of Neuronal Ion Channels in Dorsal Root Ganglion in Paclitaxel-induced Peripheral Neuropathy. Anesthesiology 120, 1463–1475 (2014).

117. Wing C, Komatsu M, Delaney SM, Krause M, Wheeler HE, Dolan ME. Application of stem cell derived neuronal cells to evaluate neurotoxic chemotherapy. Stem Cell Research 22, 79–88 (2017).

118. Bradley WG, Lassman LP, Pearce GW, Walton JN. The neuromyopathy of vincristine in man: Clinical, electrophysiological and pathological studies. Journal of the Neurological Sciences 10, 107–131 (1970).

119. Eich F, Emser W, Sybrecht GW. [Nerve conduction velocity following high-dose vincristine in the treatment of small cell bronchial cancer]. Pneumologie 44 **Suppl 1**, 574–575 (1990).

120. Boehmerle W, Huehnchen P, Peruzzaro S, Balkaya M, Endres M. Electrophysiological, behavioral and histological characterization of paclitaxel, cisplatin, vincristine and bortezomib-induced neuropathy in C57Bl/6 mice. Scientific reports 4, 6370 (2014).

121. Tanner KD, Reichling DB, Levine JD. Nociceptor Hyper-Responsiveness during Vincristine-Induced Painful Peripheral Neuropathy in the Rat. The Journal of Neuroscience 18, 6480 (1998).

122. Kavcic M, et al. Electrophysiological Studies to Detect Peripheral Neuropathy in Children Treated With Vincristine. Journal of Pediatric Hematology/Oncology 39, (2017).

123. Durens M, et al. High-throughput screening of human induced pluripotent stem cell-derived brain organoids. Journal of Neuroscience Methods 335, 108627 (2020).

124. Ghatak S, et al. NitroSynapsin ameliorates hypersynchronous neural network activity in Alzheimer hiPSC models. Molecular psychiatry 26, 5751–5765 (2021).

125. Yokoi R, et al. Analysis of signal components & 500 Hz in brain organoids coupled to microelectrode arrays: A reliable test-bed for preclinical seizure liability assessment of drugs and screening of antiepileptic drugs. Biochemistry and Biophysics Reports 28, 101148 (2021).

126. Sharf T, et al. Functional neuronal circuitry and oscillatory dynamics in human brain organoids. Nature Communications 13, 4403 (2022).

127. Odawara A, Matsuda N, Ishibashi Y, Yokoi R, Suzuki I. Toxicological evaluation of convulsant and anticonvulsant drugs in human induced pluripotent stem cell-derived cortical neuronal networks using an MEA system. Scientific reports 8, 10416 (2018).

128. Rutecki PA, Lebeda FJ, Johnston D. 4-Aminopyridine produces epileptiform activity in hippocampus and enhances synaptic excitation and inhibition. Journal of neurophysiology 57, 1911–1924 (1987).

129. Zuchero JB. Purification and culture of dorsal root ganglion neurons. Cold Spring Harbor protocols 2014, 813–814 (2014).

130. Matsuda N, Odawara A, Katoh H, Okuyama N, Yokoi R, Suzuki I. Detection of synchronized burst firing in cultured human induced pluripotent stem cell-derived neurons using a 4-step method. Biochem Biophys Res Commun 497, 612–618 (2018).

131. Kapucu FE, Tanskanen JM, Mikkonen JE, Ylä-Outinen L, Narkilahti S, Hyttinen JA. Burst analysis tool for developing neuronal networks exhibiting highly varying action potential dynamics. Frontiers in computational neuroscience 6, 38 (2012).

132. Legéndy CR, Salcman M. Bursts and recurrences of bursts in the spike trains of spontaneously active striate cortex neurons. Journal of neurophysiology 53, 926–939 (1985).

133. Lonardoni D, Marco SD, Amin H, Maccione A, Berdondini L, Nieus T. High-density MEA recordings unveil the dynamics of bursting events in Cell Cultures. In: 2015 37th Annual International Conference of the IEEE Engineering in Medicine and Biology Society (EMBC)) (2015).

134. Frey U, et al. Cell recordings with a CMOS high-density microelectrode array. Annual International Conference of the IEEE Engineering in Medicine and Biology Society IEEE Engineering in Medicine and Biology Society Annual International Conference 2007, 167–170 (2007).

135. Gandolfo M, Maccione A, Tedesco M, Martinoia S, Berdondini L. Tracking burst patterns in hippocampal cultures with high-density CMOS-MEAs. Journal of neural engineering 7, 056001 (2010).

136. Jäckel D, Frey U, Fiscella M, Franke F, Hierlemann A. Applicability of independent component analysis on high-density microelectrode array recordings. Journal of neurophysiology 108, 334–348 (2012).

137. Müller J, Bakkum D, Hierlemann A. Sub-millisecond closed-loop feedback stimulation between arbitrary sets of individual neurons. Frontiers in Neural Circuits 6, (2013).

138. Maccione A, et al. Experimental Investigation on Spontaneously Active Hippocampal Cultures Recorded by Means of High-Density MEAs: Analysis of the Spatial Resolution Effects. Frontiers in neuroengineering 3, 4 (2010).

139. Maccione A, Garofalo M, Nieus T, Tedesco M, Berdondini L, Martinoia S. Multiscale functional connectivity estimation on low-density neuronal cultures recorded by high-density CMOS Micro Electrode Arrays. Journal of Neuroscience Methods 207, 161–171 (2012).

140. Ronchi S, et al. Single-Cell Electrical Stimulation Using CMOS-Based High-Density Microelectrode Arrays. Frontiers in neuroscience 13, 208 (2019).

141. Viswam V, et al. High-density mapping of brain slices using a large multi-functional high-density CMOS microelectrode array system. In: 2017 19th International Conference on Solid-State Sensors, Actuators and Microsystems (TRANSDUCERS)) (2017).

142. Wang L, Freedman D, Sahin M, Ünlü MS, Knepper R. Active C4 Electrodes for Local Field Potential Recording Applications. Sensors (Basel, Switzerland) 16, 198 (2016).

143. Wickham J, et al. Human Cerebrospinal Fluid Induces Neuronal Excitability Changes in Resected Human Neocortical and Hippocampal Brain Slices. Frontiers in neuroscience 14, (2020).

